# An enhanced target-enrichment bait set for Hexacorallia provides phylogenomic resolution of the staghorn corals (Acroporidae) and close relatives

**DOI:** 10.1101/2020.02.25.965517

**Authors:** Peter F. Cowman, Andrea M. Quattrini, Thomas C.L. Bridge, Gregory J. Watkins-Colwell, Nur Fadli, Mila Grinblat, Thomas E. Roberts, Catherine S. McFadden, David J. Miller, Andrew H. Baird

**Affiliations:** ARC Centre of Excellence for Coral Reef Studies, James Cook University, Townsville, QLD 4811 Australia; Harvey Mudd College, Biology Department, 1250 N. Dartmouth St., Claremont, CA 91711 USA; Department of Invertebrate Zoology, National Museum of Natural History, Smithsonian Institution, Washington DC, 20560 USA; Biodiversity and Geosciences Program, Museum of Tropical Queensland, Queensland Museum, Townsville, QLD 4810, Australia; Division of Vertebrate Zoology, Yale Peabody Museum of Natural History, 170 Whitney Avenue, New Haven, Connecticut 06520, USA; Marine Science Department, Faculty of Science, Syiah Kuala University, Banda Aceh, Aceh, Indonesia; Molecular & Cell biology, College of Public Health, Medical & Vet Sciences, James Cook University, Townsville, QLD 4811 Australia

**Keywords:** Scleractinia, targeted enrichment, ultraconserved elements, UCEs, exon, phylogenetics

## Abstract

The phylogenetic utility of targeted enrichment methods has been demonstrated in taxa that often have a history of single gene marker development. These genomic capture methods are now being applied to resolve evolutionary relationships from deep to shallow timescales in clades that were previously deficient in molecular marker development and lacking robust morphological characters that reflect evolutionary relationships. Effectively capturing 1000s of loci, however, in a diverse group across a broad time scale requires a bait set that incorporates multiple baits per locus. We redesigned a custom bait set for the cnidarian class Anthozoa to target 1,436 UCE loci and 1,572 exon regions within the subclass Hexacorallia. We test this redesigned bait set on 99 specimens of hard corals (Scleractinia) spanning both the “complex” (Acroporidae, Agariciidae) and “robust” (Fungiidae) clades. With focused sampling in the staghorn coral genus *Acropora* we explore the ability of capture data to inform the taxonomy of a clade deficient in molecular resolution. A mean of 1850 (± 298) loci were captured per taxon (955 UCEs, 894 exons). A 75% complete concatenated alignment included 1792 loci (991 UCE, 801 exons) and ∼1.87 million base pairs. Parsimony informative sites varied from 48% for alignments including all three families, to 1.5% among samples within a single *Acropora* species. Maximum likelihood and Bayesian analyses recover highly resolved topologies and robust molecular relationships not previously found with traditional markers within the Acroporidae. Species level relationships within the *Acropora* genus do not support traditional morphological groups or morphological phylogenies. Both UCE and exon datasets delineated six well-supported clades within *Acropora.* The enhanced bait set for Hexacorallia will allow researchers to survey the evolutionary history of important groups of reef building corals where previous molecular marker development has been unsuccessful.

## 1. Introduction

Molecular systematics has progressed at an uneven pace across the tree of life. Several plant and animal branches on the tree of life have benefited the most from the development of single gene mitochondrial and nuclear markers resulting in large scale phylogenies (Jetz et al., 2012; Rabosky et al., 2018; Tonini et al., 2016; Upham et al., 2019) leveraging decades worth of molecular data from public resources (https://www.ncbi.nlm.nih.gov/). These large-scale phylogenies have provided a deep time perspective, systematic resolution, and often very complete biodiversity inventories of taxa. While few if any higher-level taxa can be considered complete from a species level molecular perspective, the enhanced taxonomic framework provided by these ‘Tree of Life’ projects has allowed researchers to complete shallow clades with species inventories using taxonomy-based grafting and polytomy resolution methods (Rabosky et al., 2018; Thomas et al., 2013) to incorporate species not yet sampled by genetic methods. These new methods have enabled testing of broad evolutionary and ecological hypotheses across broad taxonomic groups (Jetz et al., 2012; Pyron et al., 2013; Rabosky et al., 2018). While these approaches come with their own hurdles (Rabosky, 2015), new phylogenomic tools are facilitating a shift from the ‘top-down’ approach of sampling higher taxonomic ranks to a ‘bottom-up’ approach attempting to sample complete clades of species and even populations on shallow evolutionary time scales, leveraging high throughput sequencing technologies (Derkarabetian et al., 2018; Manthey et al., 2016; Smith et al., 2014).

Two popular phylogenomic methodologies are (i) Restriction site-associated DNA sequencing (e.g. RADseq), and (ii) Targeted sequence capture of conserved loci (Faircloth et al., 2012; Lemmon et al., 2012). Both methodologies use high-throughput sequencing, genome reduction and sample multiplexing to generate genomic datasets for hundreds of samples in substantially less time than traditional Sanger sequencing approaches (Branstetter et al., 2017). While targeted capture and RADseq data produce similar topologies (Collins and Hrbek, 2018), targeted capture of ultraconserved elements (UCEs) or exons has grown in popularity due to their comparability across taxa, tolerance of lower quality DNA templates and ability to resolve relationships at deep and shallow time scales (Faircloth et al., 2012; Harvey et al., 2016).

Targeted capture techniques were recently used for phylogenomic reconstruction across the anthozoan tree of life (Quattrini et al., 2018), with available anthozoan genomes and transcriptomes used to design a bait set to capture both UCEs and conserved exons. An average of 638 ± 222 UCE/exon loci were captured per sample, but capture rates differed greatly between Hexacorallia and Octocorallia, two major subclasses of Anthozoa, with higher recovery from the Octocorallia (soft corals, sea fans) compared with the Hexacorallia (stony corals, black corals, anemones, zoanthids). This difference in target efficiency is likely the result of the addition of octocoral-specific baits and the removal of paralogous (mostly hexacoral-specific) baits in the bait design process (Quattrini et al., 2018). In this case, separate octocoral and hexacoral-specific bait sets will increase capture efficiency and thus phylogenetic resolution across evolutionary time scales.

Within the Hexacorallia, the order Scleractinia (stony corals) has historically received significant research interest due to their role as the key ecosystem engineers on coral reefs, which host an estimated 830,000 multicellular species (Fisher et al., 2015). Hermatypic (reef-building) Scleractinia are generally colonial and represent approximately half of all scleractinian species. The capacity of stony corals to build reefs is attributed to their photosymbiotic relationship with dinoflagellates of the family Symbiodiniaceae, which mainly restricts both hermatypic corals and coral reefs to shallow tropical and sub-tropical regions (Kleypas et al., 1999). In recent years, molecular phylogenetics has fundamentally altered our understanding of the systematics and evolution of the Scleractinia, revealing that most morphological characters traditionally used to identify families, genera and species do not reflect their evolutionary history (Fukami et al., 2008, 2004; Romano and Palumbi, 1996). This has led to taxonomic revisions of the Scleractinia at every taxonomic level (Kitahara et al., 2016).

Within Scleractinia, the family Acroporidae is the most speciose family, accounting for approximately one-third of all reef-building coral species (Madin et al., 2016). Despite their ecological importance and use as a model system to understand coral biology and symbioses, there is, as yet, no well-resolved species-level molecular phylogeny for the family Acroporidae. Like plants, corals and other Anthozoans have mitochondrial substitution rates that are slower than rates of substitution across the nuclear genome (Fukami et al., 2000; Hellberg, 2006; Shearer et al., 2002; Van Oppen et al., 1999) limiting the utility of mitochondrial makers in DNA barcoding, phylogeography and shallow species-level phylogenetics in corals (Chen et al., 2009; Hellberg, 2006; Huang et al., 2008; McFadden et al., 2011). In particular, the diverse and ecologically dominant genus *Acropora* (staghorn corals) is notorious for its lack of systematic resolution with traditional mitochondrial and nuclear marker development (Chen et al., 2009). The relatively recent species level divergence and population level expansion in *Acropora* (Bellwood et al., 2017; Renema et al., 2016), means traditional mitochondrial markers offer little information to define species boundaries and robust systematic relationships. The lack of reliable molecular markers for the genus also means that the rampant synonymization of nominal species is based entirely on qualitative morphological characters (Veron and Wallace, 1984; Wallace, 1999; Wallace et al., 2012), despite their proven unreliability in determining evolutionary relationships across numerous taxonomic levels. As a result, while the genus *Acropora* contains 413 nominal species (Hoeksema and Cairns, 2019), the most recent taxonomic revision of the genus, based on morphology, recognises only 122 species (Wallace et al., 2012). Targeted sequence capture offers a new molecular tool for disentangling the relationships in hexacorals, from deep relationship among families and genera to shallow level species relationships. The family Acroporidae offers an extreme test case for the utility of a universal Hexacoral bait set for targeted sequence capture.

The aim of this study is to redesign the previously developed anthozoan UCE/exon bait set (Quattrini et al., 2018) in order to capture additional loci and enhance the capture efficiency of loci within the subclass Hexacorallia. We highlight a method for redesigning a class-level custom bait set to increase the number of captured loci within shallower clades. We use these methods to provide an enhanced RNA bait set for the anthozoan subclass Hexacorallia that targets both ultraconserved loci and exonic regions for phylogenomic reconstruction. To test the efficiency of this bait set we focus on the phylogenetically unresolved staghorn coral family Acroporidae. We include specimens from several defined morphological groups (Wallace, 1999) sampled across the Indo-Pacific with taxonomic identity determined by morphological comparisons with type materials and the original descriptions of all nominal species. We highlight the utility of targeted capture approaches for taxa that have not benefited from decades of single gene marker development. Our results demonstrate the utility of targeted capture approaches in unravelling relationships in what have been phylogenetically challenging taxa.

## 2. Materials and Methods

### 2.1. Bait Design

We re-designed a bait set, originally aimed for target-capturing UCE and exon loci in anthozoans (anthozoa-v1, Quattrini et al., 2018), to have higher specificity for and target additional loci in hexacorals. Results generated from target-capture of UCE and exon loci in 235 anthozoans (Quattrini and McFadden, unpubl. data) with the anthozoa-v1 bait set were screened to remove baits that performed poorly in hexacorals or in anthozoans more generally. Thus, 5844 baits targeting 1,123 loci (553 UCE and 570 exon loci) were retained from the anthozoa-v1 bait set. Using the program Phyluce (Faircloth, 2016) and following methods in Quattrini et al. (2018), we added additional hexacoral-specific baits to 1) improve target-capture performance of the anthozoa-v1 loci and 2) target additional hexacoral loci not included in the anthozoa-v1 bait set. Methods are briefly outlined below, however, for more details see both Quattrini et al. (2018) and the Phyluce documentation (https://phyluce.readthedocs.io/en/latest/tutorial-four.html).

To redesign baits targeting UCE loci, we first mapped 100 bp simulated-reads from the genomes of four exemplar taxa, *Acropora tenuis, Montastraea cavernosa, Amplexidiscus fenestrafer* and *Discosoma sp.*, to a masked *Nematostella vectensis (nemve)* genome (Suppl Table S1). Reads were mapped, with 0.05 substitution rate, using stampy v. 1 (Lunter and Goodson, 2011), resulting in 0.8 to 1.0% of reads aligning to the *nemve* genome. Any alignments that included masked regions (>25%) or ambiguous bases (N or X) or were too short (<80 bp) were removed using *phyluce_probe_strip_masked_loci_from_set*. An SQLite table that included regions of conserved sequences shared between *nemve* and each of the exemplar taxa was created using *phyluce_probe_get_multi_merge_table*. This table was queried using *phyluce_probe_query_multi_merge_table* to output a file containing conserved loci found in *nemve* and all other exemplar taxa.

**Table 1.** List of specimens used in the in vitro test of enhanced Hexacoral bait set with assembly summary statistics. Greyed out samples were removed from further analyses due to low contig number and/or matches to loci. Open nomenclature qualifiers (*cf., aff., sp.*) are used in species names to denote confidence in specimen identification to nominal species.

| Family | Species | Specimen No. | # Contigs | # UCE | UCE Mean Length<br>(bp) | # Exon | Exon Mean Length (bp) | Total # Loci |
| --- | --- | --- | --- | --- | --- | --- | --- | --- |
| Acroporidae | *Acropora abrotanoides* | 16-5870 | 10335 | 998 | 980 | 951 | 1075 | 1949 |
| Acroporidae | *Acropora aff. aculeus* | PN47 | 7971 | 1021 | 988 | 969 | 1069 | 1990 |
| Acroporidae | *Acropora aff. cytherea* | 74-4537 | 7393 | 1010 | 1001 | 957 | 1098 | 1967 |
| Acroporidae | *Acropora aff. cytherea* | 30-5227 | 11194 | 1000 | 1002 | 955 | 1095 | 1955 |
| Acroporidae | *Acropora aff. divaricata* | 73-3798 | 7680 | 1000 | 1041 | 944 | 1136 | 1944 |
| Acroporidae | *Acropora aff. echinata* | KM27 | 8079 | 1051 | 844 | 969 | 893 | 2020 |
| Acroporidae | *Acropora aff. echinata* | KM32 | 7965 | 1022 | 949 | 987 | 1022 | 2009 |
| Acroporidae | *Acropora aff. globiceps* | 17-6445 | 5767 | 1029 | 997 | 977 | 1082 | 2006 |
| Acroporidae | *Acropora aff. intermedia* | 30-5271 | 7164 | 986 | 1083 | 948 | 1167 | 1934 |
| Acroporidae | *Acropora aff. intermedia* | B6 | 7762 | 990 | 1003 | 933 | 1104 | 1923 |
| Acroporidae | *Acropora aff. jacquelinae* | PN54 | 6531 | 983 | 1056 | 951 | 1143 | 1934 |
| Acroporidae | *Acropora aff. latistella* | 75-4779 | 9676 | 993 | 1035 | 980 | 1112 | 1973 |
| Acroporidae | *Acropora aff. listeri* | 75-4879 | 10598 | 1010 | 819 | 945 | 866 | 1955 |
| Acroporidae | *Acropora aff. lokani* | PN51 | 10063 | 978 | 1025 | 916 | 1105 | 1894 |
| Acroporidae | *Acropora aff. longicyathus* | TER_36 | 9322 | 1021 | 782 | 966 | 830 | 1987 |
| Acroporidae | *Acropora aff. pichoni* | TER_20 | 9475 | 931 | 981 | 940 | 1095 | 1871 |
| Acroporidae | *Acropora aff. speciosa* | PN62 | 8978 | 948 | 1065 | 938 | 1153 | 1886 |
| Acroporidae | *Acropora aff. turaki* | KM35 | 8492 | 984 | 1099 | 928 | 1170 | 1912 |
| Acroporidae | *Acropora aff. turaki* | KM50 | 8194 | 1008 | 933 | 946 | 1013 | 1954 |
| Acroporidae | *Acropora azurea* | 75-4857 | 8498 | 962 | 1055 | 925 | 1154 | 1887 |
| Acroporidae | *Acropora cf. austera* | 75-4732 | 8289 | 975 | 1090 | 987 | 1183 | 1962 |
| Acroporidae | *Acropora cf. bifurcata* | 29-8193 | 9438 | 1017 | 967 | 953 | 1053 | 1970 |
| Acroporidae | *Acropora cf. digitifera* | 81-2705 | 6123 | 1002 | 940 | 947 | 1014 | 1949 |
| Acroporidae | *Acropora cf. kenti* | 75-4865 | 5133 | 1046 | 722 | 975 | 747 | 2021 |
| Acroporidae | *Acropora cf. orbicularis* | 16-5974 | 13383 | 1022 | 885 | 957 | 965 | 1979 |
| Acroporidae | *Acropora cf. orbicularis* | 29-8212 | 14180 | 984 | 1043 | 979 | 1102 | 1963 |
| Acroporidae | *Acropora cf. paniculata* | 75-4686 | 5743 | 1023 | 920 | 970 | 1005 | 1993 |
| Acroporidae | *Acropora cf. plumosa* | KM51 | 12294 | 1013 | 974 | 952 | 1067 | 1965 |
| Acroporidae | *Acropora cf. plumosa* | 69-4038 | 6009 | 993 | 997 | 968 | 1091 | 1961 |
| Acroporidae | *Acropora cf. polymorpha* | B4 | 6351 | 1013 | 953 | 956 | 1042 | 1969 |
| Acroporidae | *Acropora cf. rayneri* | TER_44 | 10527 | 929 | 957 | 928 | 1046 | 1857 |
| Acroporidae | *Acropora cf. rongelapensis* | PN44 | 7277 | 995 | 1059 | 939 | 1181 | 1934 |
| Acroporidae | *Acropora cf. rongelapensis* | PN45 | 8232 | 1023 | 968 | 969 | 1050 | 1992 |
| Acroporidae | *Acropora cf. roseni* | 29-8208 | 10316 | 992 | 1078 | 967 | 1192 | 1959 |
| Acroporidae | *Acropora cf. spicifera* | 29-8257 | 13415 | 1024 | 943 | 955 | 1042 | 1979 |
| Acroporidae | *Acropora cf. sukarnoi* | 29-8294 | 6943 | 1008 | 984 | 953 | 1079 | 1961 |
| Acroporidae | *Acropora cf. tenella* | 69-3957 | 6206 | 1004 | 1060 | 964 | 1168 | 1968 |
| Acroporidae | *Acropora cf. tenella* | PN43 | 6889 | 1024 | 923 | 995 | 1008 | 2019 |
| Acroporidae | *Acropora cf. tenella* | PN52 | 47 | 14 | 247 | 15 | 257 | 29 |
| Acroporidae | *Acropora convexa* | 30-5251 | 7817 | 1003 | 993 | 955 | 1081 | 1958 |
| Acroporidae | *Acropora kirstyae* | 74-4386 | 6885 | 995 | 1059 | 972 | 1159 | 1967 |
| Acroporidae | *Acropora lokani* | TER_8 | 12433 | 956 | 1036 | 910 | 1120 | 1866 |
| Acroporidae | *Acropora lokani* | 69-4073 | 14983 | 1012 | 948 | 919 | 1007 | 1931 |
| Acroporidae | *Acropora pichoni* | 69-3948 | 7471 | 1010 | 977 | 975 | 1063 | 1985 |
| Acroporidae | *Acropora pichoni* | TER_42 | 13089 | 995 | 981 | 942 | 1055 | 1937 |
| Acroporidae | *Acropora sarmentosa* | 74-4500 | 6377 | 1012 | 974 | 958 | 1041 | 1970 |
| Acroporidae | *Acropora solitaryensis* | 79-3886 | 10498 | 1002 | 920 | 960 | 1009 | 1962 |
| Acroporidae | *Acropora sp. (29-8273)* | 29-8273 | 7395 | 997 | 930 | 945 | 1024 | 1942 |
| Acroporidae | *Acropora sp. (29-8277)* | 29-8277 | 16145 | 1015 | 966 | 957 | 1052 | 1972 |
| Acroporidae | *Acropora sp. (30-1740)* | 30-1740 | 10037 | 1014 | 903 | 945 | 965 | 1959 |
| Acroporidae | *Acropora sp. (69-3961)* | 69-3961 | 7763 | 974 | 1034 | 953 | 1144 | 1927 |
| Acroporidae | *Acropora sp. (69-KM21)* | KM21 | 3193 | 1051 | 541 | 999 | 528 | 2050 |
| Acroporidae | *Acropora sp. (69-TER11)* | TER_11 | 14622 | 1040 | 698 | 971 | 733 | 2011 |
| Acroporidae | *Acropora sp. (69-TER35)* | TER_35 | 1498 | 657 | 297 | 521 | 287 | 1178 |
| Acroporidae | *Acropora sp. (69-TER37)* | TER_37 | 3041 | 1016 | 533 | 970 | 518 | 1986 |
| Acroporidae | *Acropora sp. (69-TER40)* | TER_40 | 11472 | 905 | 832 | 864 | 893 | 1769 |
| Acroporidae | *Acropora sp. (69-TER47)* | TER_47 | 4805 | 1051 | 833 | 999 | 887 | 2050 |
| Acroporidae | *Acropora sp. (75-4774)* | 75-4774 | 9197 | 996 | 980 | 953 | 1080 | 1949 |
| Acroporidae | *Acropora sp. (75-4803)* | 75-4803 | 3251 | 1042 | 614 | 1009 | 612 | 2051 |
| Acroporidae | *Acropora spicifera* | 30-5267 | 10635 | 993 | 1047 | 934 | 1155 | 1927 |
| Acroporidae | *Acropora syringodes* | 74-4532 | 11115 | 1007 | 1019 | 961 | 1107 | 1968 |
| Acroporidae | *Acropora vasiformis* | 16-5867 | 7941 | 1011 | 923 | 957 | 1021 | 1968 |
| Acroporidae | *Acropora verweyi* | 75-4990 | 9515 | 1003 | 1032 | 974 | 1114 | 1977 |
| Acroporidae | *Acropora walindii* | KM30 | 8091 | 1019 | 989 | 963 | 1080 | 1982 |
| Acroporidae | *Acropora walindii* | 69-4031 | 8058 | 1018 | 985 | 952 | 1077 | 1970 |
| Acroporidae | *Acropora walindii* | TER_13 | 3883 | 1045 | 817 | 991 | 880 | 2036 |
| Acroporidae | *Anacropora cf. spinosa* | 53-7072 | 43708 | 1016 | 955 | 924 | 903 | 1940 |
| Acroporidae | *Astreopora cf. gracilis* | 53-6964 | 48967 | 995 | 1011 | 920 | 984 | 1915 |
| Acroporidae | *Astreopora cf. myriophthalma* | 53-6863 | 20538 | 987 | 931 | 871 | 907 | 1858 |
| Acroporidae | *Astreopora cf. myriophthalma* | 53-7112 | 53608 | 1002 | 1034 | 936 | 1003 | 1938 |
| Acroporidae | *Astreopora cf. randalli* | 53-7115 | 49989 | 918 | 1017 | 892 | 1021 | 1810 |
| Acroporidae | *Astreopora sp. (53-7166)* | 53-7166 | 53766 | 974 | 1022 | 919 | 1027 | 1893 |
| Acroporidae | *Isopora cf. brueggemanni* | 53-7063 | 9041 | 861 | 987 | 868 | 1050 | 1729 |
| Acroporidae | *Montipora cf. aequituberculata* | 81-2751 | 11092 | 360 | 739 | 460 | 637 | 820 |
| Acroporidae | *Montipora cf. capitata* | TER_9 | 18227 | 139 | 805 | 234 | 722 | 373 |
| Acroporidae | *Montipora cf. efflorescens* | 81-2884 | 9249 | 1002 | 762 | 901 | 749 | 1903 |
| Acroporidae | *Montipora cf. foliosa* | 74-4576 | 13959 | 970 | 864 | 904 | 816 | 1874 |
| Acroporidae | *Montipora cf. monasteriata* | TER_46 | 15035 | 698 | 949 | 724 | 917 | 1422 |
| Acroporidae | *Montipora cf. nodosa* | 75-5006 | 14672 | 972 | 1001 | 913 | 963 | 1885 |
| Acroporidae | *Montipora cf. spongodes* | 81-2847 | 5295 | 1001 | 646 | 933 | 633 | 1934 |
| Acroporidae | *Montipora cf. undata* | 79-3890 | 11248 | 970 | 888 | 913 | 871 | 1883 |
| Acroporidae | *Montipora sp. (69-4301)* | 69-4301 | 15035 | 993 | 874 | 928 | 866 | 1921 |
| Acroporidae | *Montipora sp. (79-3903)* | 79-3903 | 6283 | 996 | 722 | 928 | 700 | 1924 |
| Acroporidae | *Montipora sp. (81-2697)* | 81-2791 | 7488 | 1015 | 741 | 923 | 723 | 1938 |
| Acroporidae | *Montipora sp. (81-2697)* | 81-2697 | 14906 | 984 | 846 | 884 | 837 | 1868 |
| Agariciidae | *Leptoseris aff. hawaiiensis* | PN36 | 13524 | 800 | 907 | 733 | 867 | 1533 |
| Agariciidae | *Leptoseris aff. incrustans* | PN25 | 14186 | 943 | 981 | 813 | 949 | 1756 |
| Agariciidae | *Leptoseris cf. fragilis* | PN12 | 15120 | 929 | 1038 | 811 | 983 | 1740 |
| Agariciidae | *Leptoseris cf. glabra* | PN09 | 2159 | 785 | 515 | 573 | 470 | 1358 |
| Agariciidae | *Leptoseris cf. scabra* | PN26 | 9257 | 963 | 994 | 814 | 967 | 1777 |
| Fungiidae | *Ctenactis cf. echinata* | Cf | 12759 | 977 | 987 | 814 | 926 | 1791 |
| Fungiidae | *Fungia fungites* | 60f | 15389 | 948 | 1019 | 777 | 950 | 1725 |
| Fungiidae | *Fungia fungites* | 903f | 12924 | 968 | 969 | 794 | 925 | 1762 |
| Fungiidae | *Herpolitha limax* | 4f | 15865 | 957 | 1056 | 789 | 994 | 1746 |
| Fungiidae | *Pleuractis paumotensis* | 3f | 18010 | 915 | 1038 | 762 | 975 | 1677 |
| Fungiidae | *Pleuractis paumotensis* | 11f | 14016 | 928 | 997 | 770 | 945 | 1698 |
| Fungiidae | *Polyphyllia talpina* | 7f | 15708 | 1033 | 960 | 878 | 921 | 1911 |
| Fungiidae | *Sandalolitha robusta* | 10f | 8447 | 995 | 892 | 829 | 833 | 1824 |
| Fungiidae | *Sandalolitha robusta* | 6f | 13475 | 981 | 963 | 840 | 906 | 1821 |

*Phyluce_probe_get_genome_sequences_from_bed* was used to extract these conserved regions from the *nemve* genome. Regions were then buffered to 160 bp by including an equal amount of 5’ and 3’ flanking sequence from the *nemve* genome. A temporary set of target capture baits was then designed using *phyluce_probe_get_tiled_probes*; two 120 bp baits were tiled over each locus and overlapped in the middle by 40 bp (3X density). This temporary set of baits was screened to remove baits with >25% masked bases or high (>70%) or low (<30%) GC content. At this stage, we concatenated the temporary baits with the baits retained from the anthozoa-v1 bait set and then removed any potential duplicates using-- *phyluce_probe_easy_lastz* and *phyluce_probe_remove_duplicate_hits_from_probes_using_lastz*. Bait sequences were considered duplicates if they were ≥50% identical over ≥50% of their length.

This new temporary bait set was aligned (with an identity value of 70% and a minimum coverage of 83%) to the genomes of *A. digitifera, Exaiptasia pallida, Discosoma* sp., *M. cavernosa,* and *N. vectensis* (Suppl. Table S1), and UCE loci of *Cerianthus membranaceus*, *Zoanthus* cf. *pulchellus,* and *Myriopathes ulex* (Quattrini et al., 2018) using *phyluce_probe_run_multiple_lastzs_sqlite*. From these alignments, baits that matched multiple loci were removed. We then extracted 180 bp of the sequences from the alignment files and input the data into FASTA files using *phyluce_probe_slice_sequence_from_genomes*. A list containing loci found in at least five of the taxa was created. The anthozoan UCE bait set was re-designed to target these loci using *phyluce_probe_get_tiled_probe_from_multiple_inputs*. Using this script, 120-bp baits were tiled (3X density, middle overlap) and screened for high (>70%) or low (<30%) GC content, masked bases (>25%), and duplicates as described above. Finally, the baitset was screened against the *Symbiodinium minutum* (Suppl. Table S1) genome to look for any potential symbiont loci using the scripts *phyluce_probe_run_multiple_lastzs_sqlite* and *phyluce_probe_slice_sequence_from_genomes*, with a minimum coverage of 70% and minimum identity of 60%. This UCE bait set included a total of 15,226 non-duplicated baits targeting 1,436 loci.

All of the above methods were repeated using transcriptome data to re-design the baits for target-capture of exons. We mapped 100 bp simulated-reads from the transcriptomes of three exemplar taxa, *A. digitifera,* Cerianthidae, and *Edwardsiella lineata,* to the *nemve* transcriptome (Suppl. Table S1), resulting in 4.5 to 15.3% of reads for each alignment. From these alignments, conserved sequences were added to an SQLite database. We queried this database to select loci found in *nemve* and the three exemplar taxa. Following a screening for masked regions, high/low GC content, and duplicates, a temporary exon bait set was designed. At this stage, we concatenated the temporary baits with the baits retained from the anthozoa-v1 bait set and then removed any potential duplicates using *phyluce_probe_easy_lastz* and *phyluce_probe_remove_duplicate_hits_from_probes_using_lastz*. Bait sequences were considered duplicates if they were ≥50% identical over ≥50% of their length.

The temporary baits were re-aligned to the transcriptomes of *A. digitifera,* Cerianthidae, *Protopalythoa variabilis*, *Orbicella faveolata, Pocillopora damicornis, E. lineata, N. vectensis* (Suppl. Table S1), and exon loci of *Lebrunia danae* and *Myriopathes ulex* (Quattrini et al., 2018) to ensure we could capture the loci with this bait set. We then re-designed the bait set to target these exon loci using *phyluce_probe_get_tiled_probe_from_multiple_inputs*. Using this script, 120-bp baits were tiled (3X density, middle overlap) and screened for high (>70%) or low (<30%) GC content, masked bases (>25%), and duplicates as described above. Finally, this bait set was also screened against the *S. minutum* genome to look for any potential symbiont loci using the scripts *phyluce_probe_run_multiple_lastzs_sqlite* and *phyluce_probe_slice_sequence_from_genomes*, with a minimum coverage of 70% and minimum identity of 60%. This exon bait set included a total of 15,750 non-duplicated baits targeting 1,572 loci.

The exon and UCE bait sets were concatenated and then screened to remove redundant baits (≥50% identical over >50% of their length), allowing us to create a final non-duplicated hexacoral-v2 bait set. For this study, we subset this hexacoral bait set to include baits designed against all scleractinians as well as the corallimorpharians, the antipatharian, and *N. vectensis.* This reduced the cost of the bait synthesis, while still allowing us to target a maximum number of loci. Baits were synthesized by Arbor BioSciences (Ann Arbor, MI).

### 2.2. Sample collection and morphology assessment

We tested the efficiency of this enhanced bait set on capturing loci from 99 specimens from the families Acroporidae (n=86), Agariciidae (n=5), and Fungiidae (n=9). Specimens were collected on snorkel and scuba using chisel and hammer from 2015 to 2017. Specimens spanning five of the six recognised genera of Acroporidae (*Alveopora* is unsampled) were sampled from locations across the Indo-Pacific (Suppl Table S2). The outgroup species chosen include specimens from the Agariciidae (*Leptoseris*), a close sister family of Acroporidae, and species from the more distantly related Fungiidae. Both Acroporidae and Agariciidae are members of the “complex” clade within Scleractinia, while Fungiidae is from the “robust” clade (Romano and Palumbi, 1996). This taxon sampling enables testing of the capture efficiency of the bait set at both deep (among major scleractinian clades and families) and shallow (among acroporid genera and species) evolutionary time scales.

**Table 2.** Alignment matrix statistics for different taxonomic datasets. Matrix percentage equals the percent occupancy of species per locus. PI = parsimony informative sites as calculated in Phyluce.

| **Dataset** | **#**<br>**Taxa** | **%**<br>**Matrix** | **Total #**<br>**Loci** | **# UCE/exon**<br>**Loci** | **Alignment**<br>**Length** | **Mean Aligned Locus**<br>**Length (± SD bp)** | **Aligned Locus**<br>**Length Range (bp)** | **# PI**<br>**Sites** | **% PI Sites<sup>a</sup>** |
| --- | --- | --- | --- | --- | --- | --- | --- | --- | --- |
| All Taxa | 96 | 75 | 1792 | 991/801 | 1,868,953 | 1042 ± 529 | 106-4220 | 697,843 | 46 |
| | 96 | 95 | 618 | 470/148 | 611,631 | 989 ± 413 | 222-3653 | 238,569 | 45 |
| Acroporidae | 83 | 75 | 1852 | 992/860 | 2,236,868 | 1208 ± 532 | 267-4316 | 671,600 | 38 |
| | 83 | 95 | 899 | 596/303 | 1,055,570 | 1174 ± 484 | 423-4213 | 325,959 | 38 |
| *Acropora* | 64 | 75 | 1916 | 993/923 | 2,280,802 | 1190 ± 456 | 342-3837 | 327,541 | 18 |
| | 64 | 95 | 1227 | 739/488 | 1,410,701 | 1157 ± 402 | 494-3365 | 189,608 | 16 |
| *A. walindii* | 3 | 100 | 1781 | 949/832 | 1,746,225 | 980 ± 256 | 338-3238 | 27,248<sup>b</sup> | 1.5 |

<sup>a</sup> calculated by dividing # PI sites by # sites checked for differences
<sup>b</sup> calculated as number of variable sites present in all loci

Acroporid specimens were identified by comparing skeletons to the ‘type’ material and the original descriptions of all nominal species. Uncertainties in the identifications are indicated by the use of a series of open nomenclature (ON) qualifiers (Bengtson, 1988; Sigovini et al., 2016) which provides flexibility to assign specimens to nominal species with varying degrees of certainty. Specimens with skeletons that closely resemble the original type specimen and were collected from the type location (e.g. *Acropora pichoni;* Fig. 1, Suppl Table S2) were designated as ‘topotypes’ and are given the nominal species name with no qualifier. Specimens that closely resemble the type of a nominal species but were not sampled from the type locality are given the qualifier *cf.* (e.g. *Acropora cf. plumosa;* Fig. 1,

**Figure 1.**
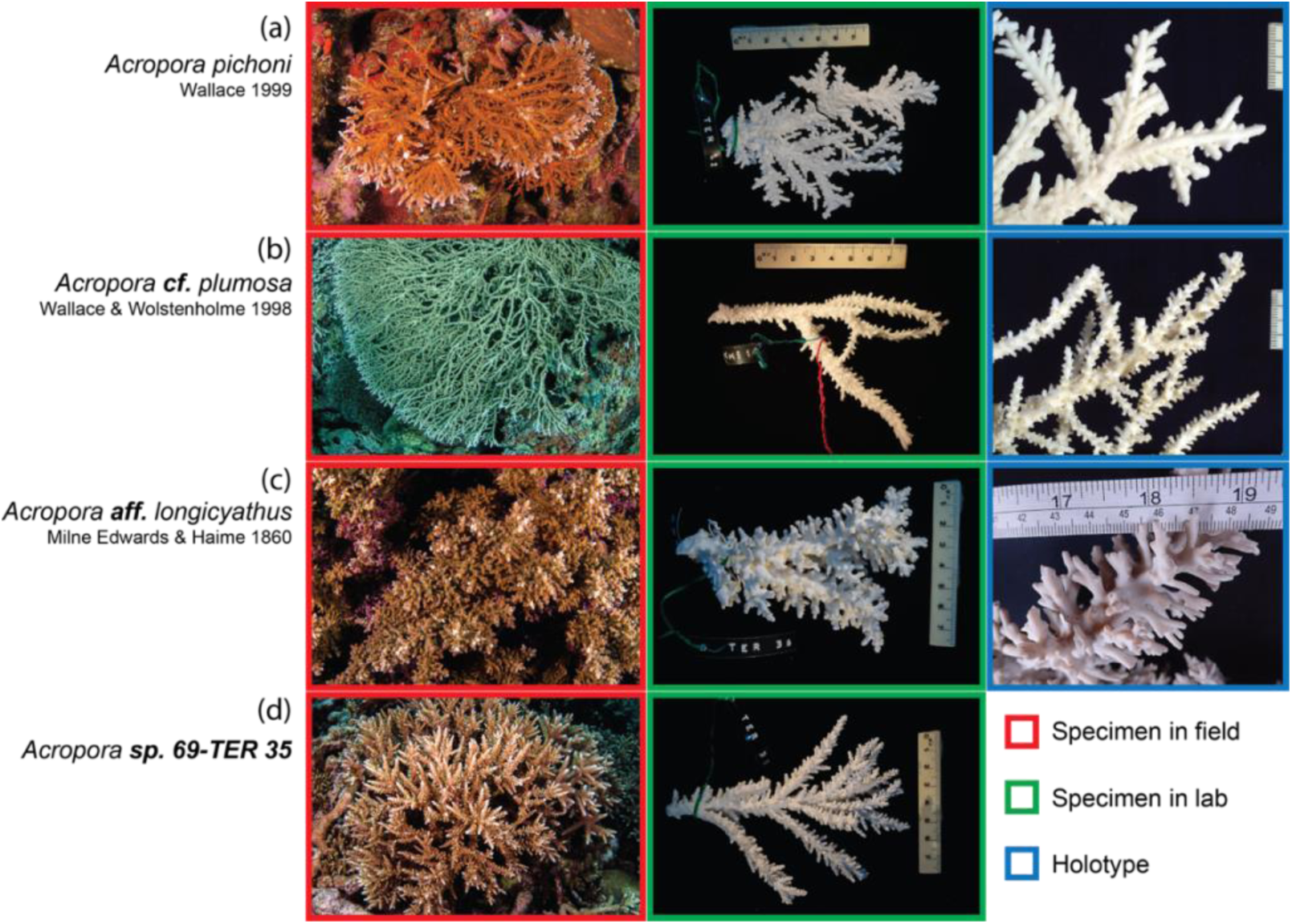
Coral specimens were identified by comparison to type material when available (*in situ* examination at MTQ; images and original descriptions). Based on these comparisons specimens were (a) identified to nominal species if they were considered a topotype sample; (b) assigned the ON qualifier *“cf”* if they resembled the type material but were not collected from the type location; (c) assigned the ON qualifier *“aff.”* if they had morphological affinities to a nominal species but could not be reliably identified using the information available; and (d) assigned the ON qualifier *“sp.”* if they showed little resemblance to any nominal species type specimen.

Suppl Table S2). Specimens that have morphological affinities to a nominal species are given the qualifier *aff.* (e.g. *Acropora aff. turaki;* Fig. 1, Suppl Table S2). These specimens may represent geographical variants of species with high morphological plasticity or undescribed species. Species that could not be matched with the type material of any nominal species were labelled as *sp.* in addition to the voucher number in its specific epithet (e.g. *Acropora sp. 69-TER 35;* Fig. 1, Suppl Table S2); these specimens are most probably undescribed species. In addition to identifying specimens with comparisons to nominal type material, we also categorized specimens into morphological grouping (“morpho-groups”) delineated by Wallace (1999) in a cladistic analysis of morphological traits. Wallace (1999) placed species into morpho-groups base on a phylogeny reconstructed from qualitative trait data. For example, the species *A. walindii* was placed in the ‘*elegans group’.* By categorizing our *Acropora* and *Isopora* specimens in this manner we can assess the molecular support for these morphological groups. Our sampling contains representatives of 16 of the 25 morpho-groups (including *Isopora)* delineated by Wallace (1999).

### 2.3. DNA extraction and target enrichment

DNA was extracted using a Qiagen DNeasy Blood & Tissue kit or a CTAB extraction protocol. DNA quality was assessed using a Nanodrop spectrophotometer, with 260/280 ratios ranging from 1.8-2.1 and 260/230 ratios ranging from 1.4-3.2. The initial concentration of each sample was measured with a Qubit 2.0 fluorometer and sent to Arbor Biosciences (Ann Arbor, MI) for library preparation.

Library preparation was performed by Arbor Biosciences following details in Quattrini et al., 2018). A total of 600 ng DNA (10 ng per μL) was sheared to a target size range of 400-800 bp using sonication, and the Kapa Hyper Prep (Kapa Biosystems) protocol was used. Universal Y-yoke oligonucleotide adapters and custom iTru dual-indexed primers as described by Glenn et al., (2019) were used. For target enrichment, the Arbor Biosciences MyBaits v. IV protocol was followed. Target enrichment was performed on pools of up to eight samples. Following target-capture enrichment, target-enriched libraries were sequenced on one lane of Illumina HiSeq 3000 (150bp PE reads). Library preparation and target enrichment were conducted by Arbor Biosciences.

### 2.4. Post-sequencing analyses

De-multiplexed Illumina reads were processed using *Phyluce* (Faircloth, 2016; http://phyluce.readthedocs.io/en/latest/tutorial-one.html/), with a few modifications. Reads were trimmed using the Illumiprocessor wrapper program (Faircloth et al., 2012) for *trimmomatic* (Bolger et al., 2014) with default values and then assembled using Spades v. 3.10 (Bankevich et al., 2012; Nurk et al., 2013). UCE and exon bait sequences were then separately matched to the assembled contigs at 70% identity and 70% coverage using *phyluce_assembly_match_contigs_to_probes*. *Phyluce_assembly_get_match_counts* and *phyluce_assembly_get_fastas_from_match_counts* were used to extract loci into FASTA files. Locus coverage was estimated using *phyluce_assembly_get_trinity_coverage* and *phyluce_assembly_get_trinity_coverage_for_uce_loci*. *Phyluce_align_seqcap_align* was used to align (with MAFFT; Katoh et al., 2002) and edge trim the loci across individuals, with default settings. Alignment matrices were created in which each locus was represented by either 75% or 95% species occupancy. Concatenated locus alignments consisted of exon loci only, UCE loci only, and all loci. The total number of variable sites, total number of parsimony informative sites, and number of parsimony informative sites per locus were calculated (using *phyluce_align_get_informative_sites*) for alignments across the following taxonomic datasets: all taxa (n=96), Acroporidae (n=83), *Acropora* (n=65), and *Acropora walindii* (n=3).

Maximum likelihood inference was conducted on each alignment (exon loci only, UCE loci only, and all loci) for both 75% and 95% data matrices (six alignment sets) using RAxML v8 (Stamatakis, 2014). This analysis was carried out using rapid bootstrapping, which allows for a complete analysis (20 ML searches and 200 bootstrap replicates) in one step. We also conducted a Bayesian analysis (100 million generations, 35% burnin) on both taxon occupancy matrices, for all loci only, using ExaBayes (Aberer et al., 2014). An extended majority rule consensus tree was produced. A GTRGAMMA model was used in both ML and Bayesian analyses.

## 3. Results

### 3.1. Bait design

The hexa-v2 bait set for all hexacorals included 25,514 baits targeting 2,499 loci (1,132 UCE and 1,367 exon loci). The bait set subset for scleractinians included 16,263 baits designed to target a total of 2,497 loci. The UCE specific bait set consisted of 8,880 baits that targeted 1,132 loci. The exon specific bait consisted of 7,383 baits that targeted 1,365 loci. In screening the combined baits set for potential hits to the *Symbiodinium* genome, 141 loci were removed. Bait sets are included as supplemental files 1 (hexa-v2) and 2 (hexa-v2-scleractinia); baits designed against transcriptomes have “design: hexatrans” in the .fasta headers.

### 3.2. Enrichment statistics and matrix results

The total number of reads obtained from Illumina sequencing ranged from 3,206 to 10,357,494 reads per sample (Suppl Table S3). Following removal of one sample (*Acropora cf. tenella*) that failed sequencing (<4K reads), and quality and adapter trimming, a mean of 3,766,889 ± 1,516,262 SD trimmed reads per sample was retained (Tables 1). Trimmed reads were assembled into a mean of 11,858 ± 9,682 SD contigs per sample (range: 1,498 to 53,766) with a mean length of 506 ± 75 bp using SPAdes (Tables 1; Table S3). Read coverage per contig ranged from 0.3 to 75X.

A total of 1131 UCE loci and 1332 exon loci (2463 total loci out of 2,497 targeted loci) were recovered from the assembled contigs (Fig. 2, Table 1). Following the removal of two samples (*Montipora cf. aequituberculata*, *Montipora cf. capitata*) due to relatively few loci recovered (Fig. 2, <1000 loci), mean number of loci was 1,900 ± 140 SD per sample (range: 1,178 to 2,051). We recovered slightly higher numbers of loci from acroporids (1,930 ± 116 SD) than from agariciids (1,632 ± 182) and fungiids (1,773 ± 73 SD). The total number of UCE loci recovered was 983 ± 63 SD per sample (range: 657 to 1051) with a mean length of 933 ± 139 bp (range: 297 to 1,099 bp). The total number of exon loci recovered was 917 ± 82 (range: 521 to 1009) with a mean length of 976 ± 168 bp (range: 287 to 1,192 bp) (Suppl Table S3). Read coverage per locus ranged from 1 to 337X for UCE loci and 0.5 to 270X for exon loci.

**Figure 2.**
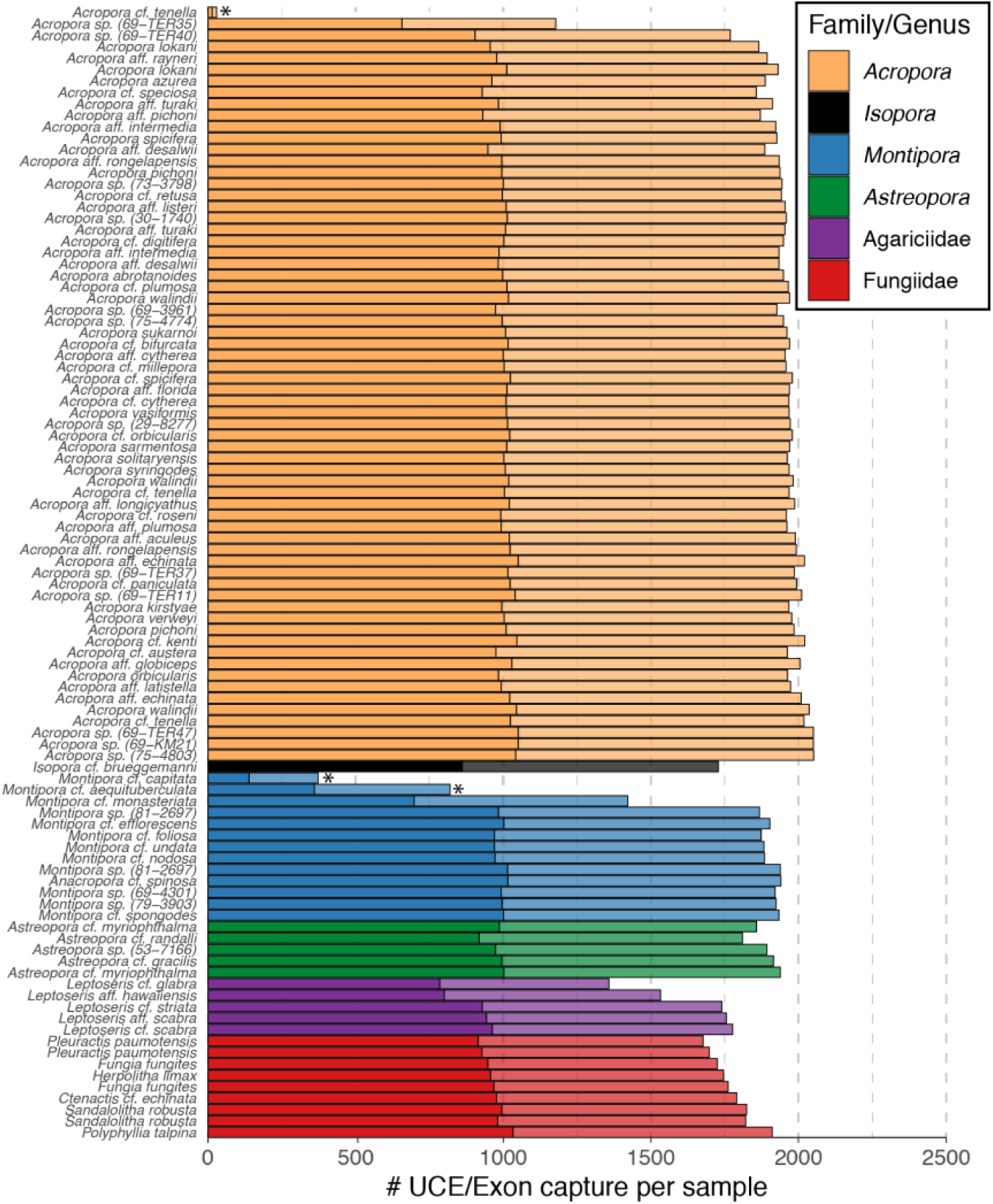
Number of UCE (solid) and exon (transparent) loci captured per taxa. Colours reflect family/genus membership. * denotes samples that were subsequently removed due to low capture rate.

Extracted loci were aligned and pruned to form 75% and 95% occupancy matrices for UCEs only and exons only. The final UCE only alignments consisted of 991 and 470 loci (75% and 95% matrices respectively) for 96 samples that passed quality assessment. The final exon only alignments contained 801 and 148 loci (75% and 95% matrices respectively) for the 96 samples. The final combined concatenated alignments contained 1792 UCE/exon loci in the 75% matrix and 618 in the 95% matrix (Table 2). The number of parsimony informative sites varied from 48% in alignments containing all three families (Acroporidae, Agariciidae, Fungiidae), to 1.5% among samples within a single *Acropora* species (*A. walindii*). Within the family Acroporidae parsimony informative sites accounted for 38% among genera, and 18% within the genus *Acropora* (Table 2).

### 3.4. Coral phylogenomics

All six alignment matrices produced robust phylogenies separating the three families and the deep split between “complex” (Acroporidae + Agariciidae) and “robust” (Fungiidae) clades within the Scleractinia (Fig. 3; Suppl Figs S1-3). Given the congruent topologies across alignment sets, we discuss all systematic relationships below referring to the combined UCE/exon data alignments with 75% completeness (Figs 3-4).

**Figure 3.**
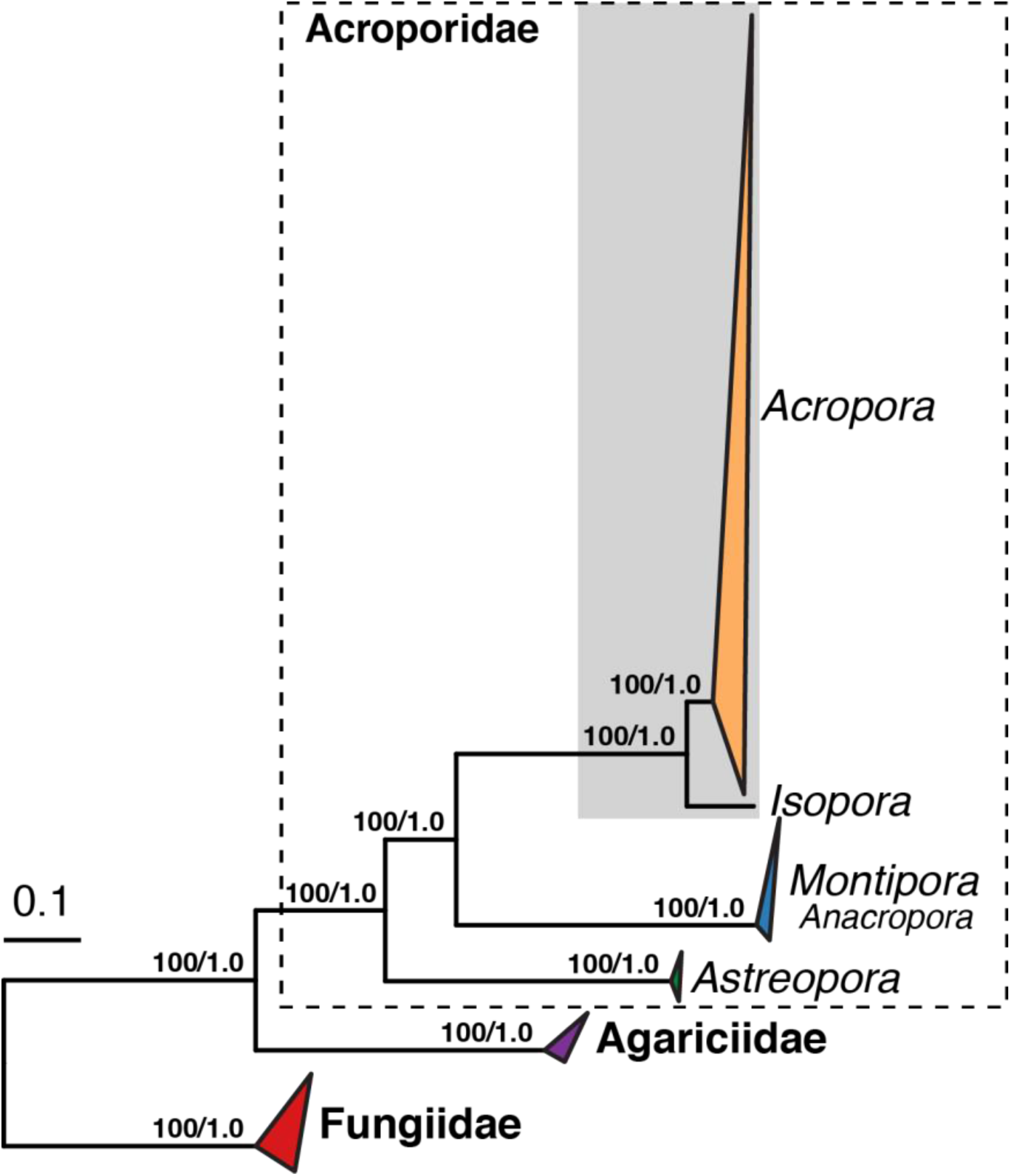
RAxML phylogeny based on 75% UCE/exon complete matrix. Phylogenetic relatiohsip are delineated between Acroporidae genera (dashed outline) and outgroup families Agariciidae and Fungiidae. Acroporidae and Agariciidae are both from the complex scleractinian clade, while Fungiidae is from the robust clade (Romano & Palumbi, 1996). Node values show RAxML bootstrap values and posterior probabilities from ExaBayes analysis. Species level relationships within the Acropora/Isopora clade (in grey box) are shown in Figure 4.

Both maximum likelihood (ML) and Bayesian analyses resulted in high support for nodes in resulting consensus topologies (Fig. 3, Suppl Fig. S4). In the ML tree 83% of internal nodes are resolved with RAxML bootstrap values of 100 (92% ≥ 70), and in the Bayesian analyses 94% of nodes received posterior probabilities of 1 (96% ≥ 0.7; Suppl Fig. S4). Within the Acroporidae, the genus *Astreopora* is the first lineage to branch off in the family, followed by *Montipora* which contains *Anacropora* (Fig. 4). *Isopora* and *Acropora* are recovered as sister clades but independent genera. Within *Acropora*, all phylogenetic reconstructions highlight six major clades with high resolution (Fig. 4). Our molecular phylogeny indicates that most of the morphological species groups of Wallace (1999) are paraphyletic (9 of the 16 groups represented) (Fig. 4 inset). For example, species assigned to the *robusta* group appear in clades VI, V and III in the phylogeny (Fig. 4). Similarly, representatives of the *echinata, aspera* and *divaricata* groups are found across multiple clades. Representatives of some morpho-groups (e.g. *elegans* and *hyacinthus* groups) occur in the same molecular clade, but are paraphyletic at the species level, and do not reflect morphological relationships among member species (Fig. 4).

**Figure 4.**
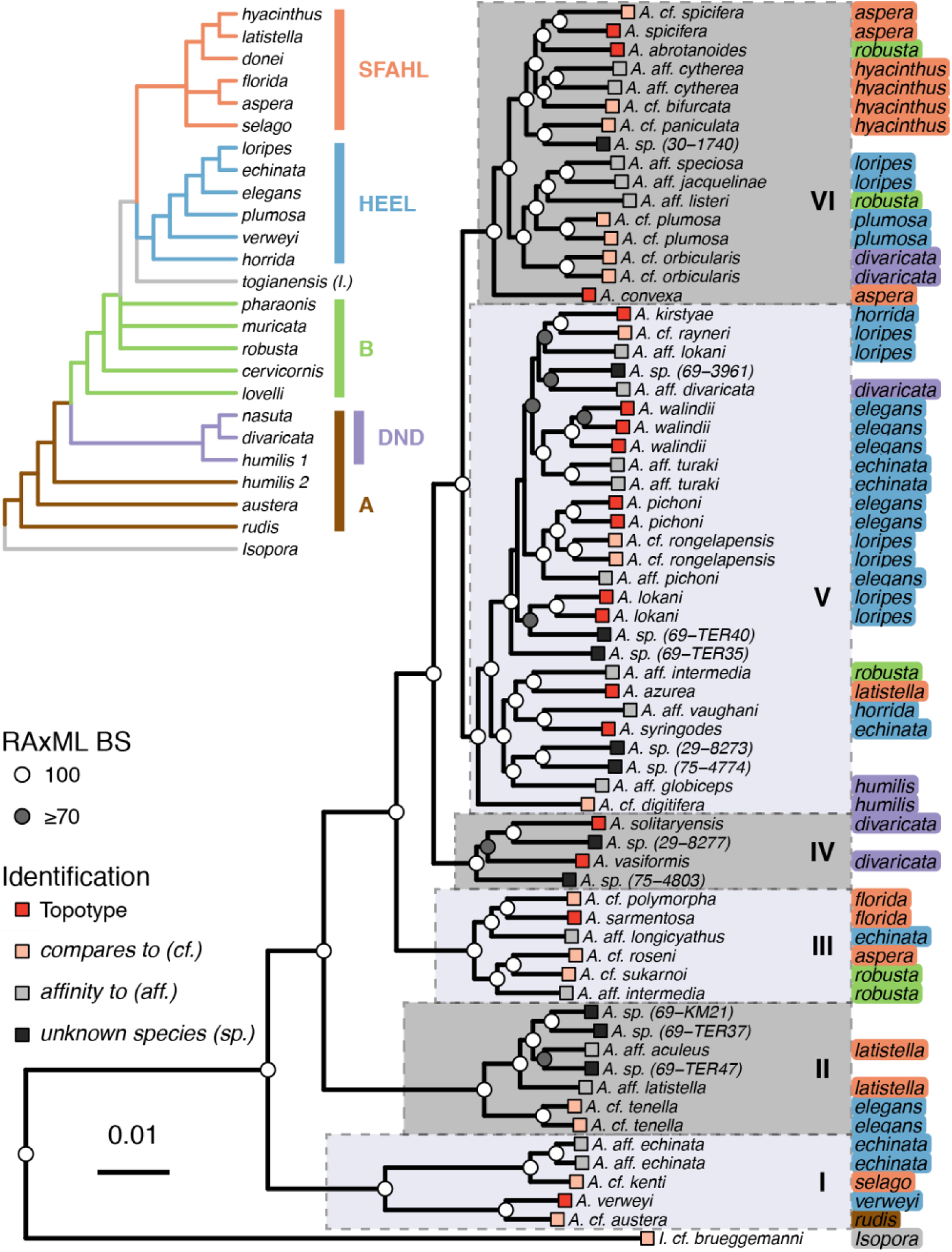
Phylogeny of *Acropora* and *Isopora* extracted from the ML analysis of the combined UCE/exon 75% complete matrix. Inset cladogram depicts the systematic relationships of morpho-groups from Wallace (1999) with branch colours highlighting the major named morphological clades: Clade A (*rudis, austera* and *humilis 2* groups) which contains the DND clade (*digitifera-nasuta-divaricata* and *humilis* group 1); clade B (*lovelli, cervicornis, robusta, muricata, pharonis*); the HEEL clade (*horrida-elegans-echinata-loripes*); the SFAHL clade (*selago-florida-aspera-hyacinthus-latistella*). Specimen are labels indicate morpho-group designation coloured by morphological clade (inset). Squares at tips reflect the ON identification for each specimen (see legend). Bootstrap values of either 100% (white) or greater than 70% (grey) are indicating on internal nodes. Robust molecular clades are delineated I-VI.

Of the 65 *Acropora* specimens, 17 were classified as topotypes and 18 designated as *cf.* In combination, these 35 specimens represent 26 nominal species of *Acropora*. In addition, 20 specimens had affinities (*aff.*) to nominal species and 12 samples could not be matched to any of the 414 nominal species (Suppl Table S1). In general, specimens given the same open nomenclature clustered together with high bootstrap support (e.g. *A. walindii, A. aff cytherea, A. cf plumosa*), as did the specimens identified as topotypes and those that matched the type from another region (e.g. *A. spicifera and A. cf spicifera*) (Fig. 4). However, not surprisingly, specimens with an affinity to a given nominal species often occurred in different clades (e.g *A. aff. pichoni, A. aff. intermedia,* Fig. 4) suggesting that the morphological characters on which these affinities were qualitatively assessed (gross morphology and radial corallite shape) are not phylogenetically informative.

## 4. Discussion

### 4.1 An enhanaced target-enrichment baits set for Hexacorallia

Here, we provide a pipeline to redesign a class level UCE/exon bait set (Anthozoa; (Quattrini et al., 2018) to improve the capture efficiency of loci from a variety of taxa within a lower taxonomic rank (subclass Hexacorallia). Our new bait set, however, also includes a subset of the originally targeted loci (anthozoa-v1), which will allow for future comparisons and integration with the original probe set. We tested our enhanced hexacoral UCE/exon probe design by focusing on a phylogenomic reconstruction of the staghorn corals (Acroporidae), while also including families from both complex (Agariciidae) and robust (Fungiidae) clades within the Scleractinia. This new enhanced probe set increases the capture efficiency of UCE/exons for scleractinians when compared to the class-level design (Quattrini et al., 2018) and results in higher phylogenetic resolution within an important group of reef building corals.

Our new bait design greatly increases the number of UCE and exon loci captured within the Hexacorallia (Fig. 2; Table 1, 2). Overall, there is an almost four-fold increase in the number of loci captured with the hexa-v2 bait set when compared to the original antho-v1 bait set. Resulting alignments retain higher numbers of loci for phylogenomic reconstruction within hexacorals in more complete alignment matrices: 1792 and 618 UCE/exon loci in 75% and 95% complete matrices respectively, compared to 438 UCE/exon loci in a 50% complete matrix (Quattrini et al., 2018). In addition, individual locus alignments are longer (1042 ± 529 bp; Table 2) when compared to those captured from the antho-v1 probe design (205 ± 93 bp; Quattrini et al., 2018), with a greater number of parsimony informative sites (46%; Table 2). Fewer numbers of reads and contigs were recovered in this study compared to Quattrini et al., (2018) therefore, it is possible that more on-target reads were obtained using this new bait set and/or the MyBaits v IV protocol, which was not used in Quattrini et al., (2018). In parallel to our redesign approach of the Anthozoa probe set, Quek et al., (2020) recently released a transcriptome-based targeted-enrichment bait set focused on Scleractinia targeting 1,139 exon regions. This Scleractinia focused exon bait set appears to have no overlap with the targeted exons of the Antho-v1 bait design (Quattrini et al., 2018; Quek et al., 2020) and only overlap in 12 exon loci, at a 50% similarity level, in our enhanced hexa-v2 scleractinia bait set (AMQ). While the dissimilarity are like due to the different taxonomic focus (Quek et al., 2020), future combinations of these independent bait designs will improve and expand the phylogenomic resolution of corals.

### 4.2 Coral systematics and taxonomy

The redesigned bait set provides a high throughput tool for phylogenetic inference in a systematically challenging group of corals. Initial molecular assessment of numerous groups within the Scleractinia have highlighted the fact that traditional morphological taxonomic schemes do not reflect systematic relationships or the evolutionary history revealed by molecular phylogenetics (Fukami et al., 2004; Huang et al., 2014; Kitahara et al., 2016; Romano and Palumbi, 1996). Similarly, our targeted capture data identifies many problems with the current systematics of the Acroporidae (Fig. 4), in particular, the incongruence between morphological and molecular based phylogenies (see also Richards et al., 2013). Unlike many other groups that have gained systematic resolution through the sequencing of single gene mitochondrial and nuclear markers, such markers provided limited phylogenetic resolution in the Scleractinia, particularly the hyperdiverse Acroporidae.

However, a handful of informative markers have been exploited in an opportunistic way for phylogenetics in some clades. For example, the mitochondrial spacer region between COI and 16S rRNA has been used to resolve species boundaries in agariciid genera *Leptoseris* and *Pavona* from Hawaii (Luck et al., 2013; Pochon et al., 2015; Terraneo et al., 2017). Similarly, the putative mitochondrial control region and an open reading frame located between the mitochondrial ATP6 and NAD4 genes have provided resolution among genera in the family Pocilloporidae (Flot et al., 2011; Pinzón et al., 2013; Schmidt-Roach et al., 2013). However, the utility of these and other markers are not universal across coral taxa and sampling has not been uniform across clades or genes.

Obtaining informative phylogenetic markers capable of delineating species has been particularly problematic in the Acroporidae (but see Van Oppen et al., 2000; Márquez, Van Oppen, Willis, Reyes, & Miller, 2002). A recent phylogenetic reconstruction for the family Acroporidae (Huang et al., 2018) which mined two decades of molecular data resulted in an alignment representing a total 119 accepted species across seven mitochondrial markers and two nuclear markers (ITS1 and 2, Pax-C). However, species sampling varied greatly among gene datasets (n = 17-73 species), resulting in a concatenated matrix that was 35% complete and where 25% of sampled species were represented by a single gene (Huang et al., 2018). Our testing of the enhanced bait set resulted in a 75% complete matrix consisting of 1792 UCE/exon loci for specimens linked to 26 nominal and 32 potentially undescribed species, where the lowest number of loci for a single species in the alignment was 976. Phylogenomic reconstructions of our UCE/exon dataset agree with other molecular phylogenetic studies of Scleractinia with a deep split between “complex” and “robust” corals (Romano and Palumbi, 1996; Ying et al., 2018) and the relationships among acroporid genera (Fig. 3; Fukami et al., 2000). At a shallower scale, this enhanced bait design provided a level of resolution within the genus *Acropora* that has not been achieved with available single markers or morphological analysis.

### 4.3. Phylogenomic resolution of staghorn coral

Phylogenomic reconstruction, using both ML and Bayesian methods, consistently resolved six molecular clades within *Acropora* with high support (bootstrap = 100; posterior =1), regardless of alignment type (Fig. 4; Suppl Figs S1-S4). While some traditional morphological characters seem to distinguish nominal species lineages they appear to offer little congruence with the molecular systematic relationships reconstructed here (Fig. 4) and in previous single gene reconstructions within *Acropora* (Huang et al., 2018; Richards et al., 2016, 2013). Richards et al. (2013) found 6-7 clades within *Acropora*, although the phylogenetic resolution was too low to support relationships among and within the clades.

Despite little overlap in the species sampled, deep splits in the single gene tree of Richards et al. (2013) show some concordance with the arrangement of Clade I and II in our reconstruction (Fig. 4.), however, other clades differ considerably. Our sampling only presents a small fraction of the diversity of *Acropora* and so increased sampling effort could reveal further differentiation within and among the molecular clades highlighted here. Given our sampling includes a broad range of traditional morpho-groupings (Fig. 4) from across the Indo-Pacific (Table S2), it is probable that the genus *Acropora* is broadly represented by the six molecular clades found here.

## 5. Conclusions

Our enhanced hexacoral bait set has the ability to generate new phylogenomic datasets to resolve deep to shallow-level evolutionary relationships among reef building corals and their relatives. This new bait set improved on the capture efficiency of the previous anthozoan bait design resulting in higher numbers of UCE and exons in more complete and longer alignments. Our subsequent phylogenomic analyses demonstrated that the macromorphological characters traditionally used for taxonomic identification in corals do not reflect evolutionary relationships. As climate change impacts coral reefs around the world, conservation efforts rely on a robust taxonomy (Thomson et al., 2018) and a phylogenetic framework in which to assess extinction risk (Huang, 2012). Importantly, over 50 % of our specimens cannot readily be assigned to any of the 414 nominal species of *Acropora*, suggesting the true diversity of this genus has been seriously underestimated in recent revisions (Wallace et al., 2012). Our new bait set, in conjunction with wider geographic sampling of species and a close examination of the type material, will provide a renewed taxonomic focus for reef building corals.

## Acknowledgements

This research was funded through the ARC DECRA to PFC (DE170100516), the ARC Centre of Excellence Programme (CE140100020) to AHB and TB, the NSF-DEB #1457817 to CSM. We thank B. Faircloth for guidance on bait design, M de Freitas Prazeras for tissue sample curation and DNA extraction. For access to type material and images we thank: P. Muir, M. Lowe, A. Cabrinovic, K. Johnson, P. Joannot, P. Lozouet, M., H. Sattmann, C. Lüter, T. Coffer, E. Lazo-Wasem, L. Rojas, M. Sørensen; D. Matenaar, W. Y. Licuanan, C. Veron, N. Raymond. For assistance in the field we thank: Nurma (Oong Bungalow’s Pulau Weh); Muslim (Pulau Beras); staff at Mahonia na Dari (Kimbe Bay) and Nihco Marine Park (Pohnpei); Lizard Island Research Station, D. Huang and A. Bauman (National University of Singapore); Jetty Dive (Coffs Harbour); staff at Lord Howe Island Marine Parks and the Lord Howe Island Board (Lord Howe Island); B. Busteed (Howea Dive, Lord Howe Island); H. McDonald (Solitary Island Marine Park).

## Author Contributions

PFC designed the project with help from AMQ and CSM. Sample collection was conducted by AB, TB, TER and MG. Morphological assessment of acroporids was conducted by TB and AB. The hexa-2 bait set was designed by AMQ. AMQ assembled contigs and performed phylogenomic analyses. Initial draft of paper was written by PFC and AMQ with input from TB. All authors contributed to subsequent drafts.

## Data Accessibility

Tree and alignment files: Data Dryad Entry https://doi.org/10.5061/dryad.9p8cz8wc8; Raw Data: SRA GenBank SUB6852542, BioSample #SAMN13871686-1781; Hexacoral bait set: Supplemental Files 2 and 3, Data Dryad Entry https://doi.org/10.5061/dryad.9p8cz8wc8

## References

1. Aberer, A.J., Kobert, K., Stamatakis, A., 2014. Exabayes: Massively parallel bayesian tree inference for the whole-genome era. Mol. Biol. Evol. 31, 2553–2556. https://doi.org/10.1093/molbev/msu236

2. Bankevich, A., Nurk, S., Antipov, D., Gurevich, A.A., Dvorkin, M., Kulikov, A.S., Lesin, V.M., Nikolenko, S.I., Pham, S., Prjibelski, A.D., Pyshkin, A. V., Sirotkin, A. V., Vyahhi, N., Tesler, G., Alekseyev, M.A., Pevzner, P.A., 2012. SPAdes: A new genome assembly algorithm and its applications to single-cell sequencing. J. Comput. Biol. 19, 455–477. https://doi.org/10.1089/cmb.2012.0021

3. Bellwood, D.R., Goatley, C.H.R., Bellwood, O., 2017. The evolution of fishes and corals on reefs: form, function and interdependence. Biol. Rev. 92, 878–901. https://doi.org/10.1111/brv.12259

4. Bengtson, P., 1988. Open Nomenclature. Palaeontology 31, 223–227.

5. Bolger, A.M., Lohse, M., Usadel, B., 2014. Trimmomatic: A flexible trimmer for Illumina sequence data. Bioinformatics 30, 2114–2120. https://doi.org/10.1093/bioinformatics/btu170

6. Branstetter, M.G., Longino, J.T., Ward, P.S., Faircloth, B.C., 2017. Enriching the ant tree of life: enhanced UCE bait set for genome-scale phylogenetics of ants and other Hymenoptera. Methods Ecol. Evol. 8, 768–776. https://doi.org/10.1111/2041-210X.12742

7. Chen, I.P., Tang, C.Y., Chiou, C.Y., Hsu, J.H., Wei, N.V., Wallace, C.C., Muir, P., Wu, H., Chen, C.A., 2009. Comparative analyses of coding and noncoding DNA regions indicate that Acropora (Anthozoa: Scleractina) possesses a similar evolutionary tempo of nuclear vs. mitochondrial genomes as in plants. Mar. Biotechnol. 11, 141–152. https://doi.org/10.1007/s10126-008-9129-2

8. Collins, R.A., Hrbek, T., 2018. An in silico comparison of protocols for dated phylogenomics. Syst. Biol. 67, 633–650. https://doi.org/10.1093/sysbio/syx089

9. Derkarabetian, S., Starrett, J., Tsurusaki, N., Ubick, D., Castillo, S., Hedin, M., 2018. A stable phylogenomic classification of Travunioidea (Arachnida, Opiliones, Laniatores) based on sequence capture of ultraconserved elements. Zookeys 760, 1–36. https://doi.org/10.3897/zookeys.760.24937

10. Faircloth, B.C., 2016. PHYLUCE is a software package for the analysis of conserved genomic loci. Bioinformatics 32, 786–788. https://doi.org/10.1093/bioinformatics/btv646

11. Faircloth, B.C., McCormack, J.E., Crawford, N.G., Harvey, M.G., Brumfield, R.T., Glenn, T.C., 2012. Ultraconserved elements anchor thousands of genetic markers spanning multiple evolutionary timescales. Syst. Biol. 61, 717–726. https://doi.org/10.1093/sysbio/sys004

12. Fisher, R., O’Leary, R.A., Low-Choy, S., Mengersen, K., Knowlton, N., Brainard, R.E., Caley, M.J., O’Leary, R.A., Low-Choy, S., Mengersen, K., Knowlton, N., Brainard, R.E., Caley, M.J., O’Leary, R.A., Low-Choy, S., Mengersen, K., Knowlton, N., Brainard, R.E., Caley, M.J., O’Leary, R.A., Low-Choy, S., Mengersen, K., Knowlton, N., Brainard, R.E., Caley, M.J., 2015. Species Richness on Coral Reefs and the Pursuit of Convergent Global Estimates. Curr. Biol. 25, 500–505. https://doi.org/10.1016/j.cub.2014.12.022

13. Flot, J.F., Blanchot, J., Charpy, L., Cruaud, C., Licuanan, W.Y., Nakano, Y., Payri, C., Tillier, S., 2011. Incongruence between morphotypes and genetically delimited species in the coral genus Stylophora: Phenotypic plasticity, morphological convergence, morphological stasis or interspecific hybridization? BMC Ecol. 11, 1–14. https://doi.org/10.1186/1472-6785-11-22

14. Fukami, H., Budd, A.F., Paulay, G., Solé-Cava, A., Chen, C.A., Iwao, K., Knowlton, N., 2004. Conventional taxonomy obscures deep divergence between Pacific and Atlantic corals. Nature 427, 832–835. https://doi.org/10.1038/nature02339

15. Fukami, H., Chen, C.A., Budd, A.F., Collins, A., Wallace, C.C., Chuang, Y.-Y., Chen, C., Dai, C.-F., Iwao, K., Sheppard, C., Knowlton, N., 2008. Mitochondrial and Nuclear Genes Suggest that Stony Corals Are Monophyletic but Most Families of Stony Corals Are Not (Order Scleractinia, Class Anthozoa, Phylum Cnidaria). PLoS One 3, e3222. https://doi.org/10.1371/journal.pone.0003222

16. Fukami, H., Omori, M., Hatta, M., 2000. Phylogenetic Relationships in the Coral Family Acroporidae, Reassessed by Inference from Mitochondrial Genes. Zoolog. Sci. 17, 689– 696. https://doi.org/10.2108/zsj.17.689

17. Glenn, T.C., Nilsen, R.A., Kieran, T.J., Sanders, J.G., Bayona-Vásquez, N.J., Finger, J.W., Pierson, T.W., Bentley, K.E., Hoffberg, S.L., Louha, S., Garcia-De Leon, F.J., Del Rio Portilla, M.A., Reed, K.D., Anderson, J.L., Meece, J.K., Aggrey, S.E., Rekaya, R., Alabady, M., Belanger, M., Winker, K., Faircloth, B.C., 2019. Adapterama I: Universal stubs and primers for 384 unique dual-indexed or 147,456 combinatorially-indexed Illumina libraries (iTru & iNext). PeerJ 2019, 1–31. https://doi.org/10.7717/peerj.7755

18. Harvey, M.G., Smith, B.T., Glenn, T.C., Faircloth, B.C., Brumfield, R.T., 2016. Sequence Capture versus Restriction Site Associated DNA Sequencing for Shallow Systematics. Syst. Biol. 65, 910–924. https://doi.org/10.1093/sysbio/syw036

19. Hellberg, M.E., 2006. No variation and low synonymous substitution rates in coral mtDNA despite high nuclear variation. BMC Evol. Biol. 6, 1–8. https://doi.org/10.1186/1471-2148-6-24

20. Hoeksema, B.W., Cairns, S.D., 2019. World List of Scleractinia [WWW Document]. URL http://www.marinespecies.org/scleractinia (accessed 12.20.19).

21. Huang, D., 2012. Threatened Reef Corals of the World. PLoS One 7, e34459. https://doi.org/10.1371/journal.pone.0034459

22. Huang, D., Benzoni, F., Fukami, H., Knowlton, N., Smith, N.D., Budd, A.F., 2014. Taxonomic classification of the reef coral families Merulinidae, Montastraeidae, and Diploastraeidae (Cnidaria: Anthozoa: Scleractinia). Zool. J. Linn. Soc. 171, 277–355. https://doi.org/10.1111/zoj.12140

23. Huang, D., Goldberg, E.E., Chou, L.M., Roy, K., 2018. The origin and evolution of coral species richness in a marine biodiversity hotspot*. Evolution (N. Y). 72, 288–302. https://doi.org/10.1111/evo.13402

24. Huang, D., Meier, R., Todd, P.A., Chou, L.M., 2008. Slow mitochondrial COI sequence evolution at the base of the metazoan tree and its implications for DNA barcoding. J. Mol. Evol. 66, 167–174. https://doi.org/10.1007/s00239-008-9069-5

25. Jetz, W., Thomas, G.H., Joy, J.B., Hartmann, K., Mooers, a O., 2012. The global diversity of birds in space and time. Nature 491, 444–448. https://doi.org/10.1038/nature11631

26. Katoh, K., Misawa, K., Kuma, K., Miyata, T., 2002. MAFFT: a novel method for rapid multiple sequence alignment based on fast Fourier transform. Nucleic Acids Res. 30, 3059–66.

27. Kitahara, M. V, Fukami, H., Benzoni, F., Huang, D., 2016. The New Systematics of Scleractinia: Integrating Molecular and Morphological Evidence, in: Goffredo, S., Dubinsky, Z. (Eds.), The Cnidaria, Past, Present and Future: The World of Medusa and Her Sisters. Springer International Publishing, Cham, pp. 41–59. https://doi.org/10.1007/978-3-319-31305-4_4

28. Kleypas, J.A., McManu, J.W., Mene, L.A.B., 1999. Environmental limits to coral reef development: Where do we draw the line? Am. Zool. 39, 146–159. https://doi.org/10.1093/icb/39.1.146

29. Lemmon, A.R., Emme, S. a, Lemmon, E.M., 2012. Anchored hybrid enrichment for massively high-throughput phylogenomics. Syst. Biol. 61, 727–744. https://doi.org/10.1093/sysbio/sys049

30. Luck, D.G., Forsman, Z.H., Toonen, R.J., Leicht, S.J., Kahng, S.E., 2013. Polyphyly and hidden species among Hawai’i’s dominant mesophotic coral genera, Leptoseris and pavona (Scleractinia: Agariciidae). PeerJ 2013, 1–20. https://doi.org/10.7717/peerj.132

31. Lunter, G., Goodson, M., 2011. Stampy: A statistical algorithm for sensitive and fast mapping of Illumina sequence reads. Genome Res. 21, 936–939. https://doi.org/10.1101/gr.111120.110

32. Madin, J.S., Hoogenboom, M.O., Connolly, S.R., Darling, E.S., Falster, D.S., Huang, D., Keith, S.A., Mizerek, T., Pandolfi, J.M., Putnam, H.M., Baird, A.H., 2016. A Trait-Based Approach to Advance Coral Reef Science. Trends Ecol. Evol. 31, 419–428. https://doi.org/10.1016/j.tree.2016.02.012

33. Manthey, J.D., Campillo, L.C., Burns, K.J., Moyle, R.G., 2016. Comparison of target-capture and restriction-site associated DNA sequencing for phylogenomics: A test in cardinalid tanagers (Aves, Genus: Piranga). Syst. Biol. 65, 640–650. https://doi.org/10.1093/sysbio/syw005

34. Márquez, L.M., Van Oppen, M.J.H., Willis, B.L., Reyes, A., Miller, D.J., 2002. The highly cross-fertile coral species, Acropora hyacinthus and Acropora cytherea, constitute statistically distinguishable lineages. Mol. Ecol. 11, 1339–1349. https://doi.org/10.1046/j.1365-294X.2002.01526.x

35. McFadden, C.S., Benayahu, Y., Pante, E., Thoma, J.N., Nevarez, P.A., France, S.C., 2011. Limitations of mitochondrial gene barcoding in Octocorallia. Mol. Ecol. Resour. 11, 19– 31. https://doi.org/10.1111/j.1755-0998.2010.02875.x

36. Nurk, S., Bankevich, A., Antipov, D., Gurevich, A.A., Korobeynikov, A., Lapidus, A., Prjibelski, A.D., Pyshkin, A., Sirotkin, A., Sirotkin, Y., Stepanauskas, R., Clingenpeel, S.R., Woyke, T., McLean, J.S., Lasken, R., Tesler, G., Alekseyev, M.A., Pevzner, P.A., 2013. Assembling single-cell genomes and mini-metagenomes from chimeric MDA products. J. Comput. Biol. 20, 714–737. https://doi.org/10.1089/cmb.2013.0084

37. Pinzón, J.H., Sampayo, E., Cox, E., Chauka, L.J., Chen, C.A., Voolstra, C.R., Lajeunesse, T.C., 2013. Blind to morphology: Genetics identifies several widespread ecologically common species and few endemics among Indo-Pacific cauliflower corals (Pocillopora, Scleractinia). J. Biogeogr. 40, 1595–1608. https://doi.org/10.1111/jbi.12110

38. Pochon, X., Forsman, Z.H., Spalding, H.L., Padilla-Gamiño, J.L., Smith, C.M., Gates, R.D., 2015. Depth specialization in mesophotic corals (Leptoseris spp.) and associated algal symbionts in Hawai‘i. R. Soc. Open Sci. 2. https://doi.org/10.1098/rsos.140351

39. Pyron, R.A., Burbrink, F.T., Wiens, J.J., 2013. A phylogeny and revised classification of Squamata, including 4161 species of lizards and snakes. BMC Evol. Biol. 13, 93. https://doi.org/10.1186/1471-2148-13-93

40. Quattrini, A.M., Faircloth, B.C., Dueñas, L.F., Bridge, T.C.L., Brugler, M.R., Calixto-Botía, I.F., DeLeo, D.M., Forêt, S., Herrera, S., Lee, S.M.Y., Miller, D.J., Prada, C., Rádis-Baptista, G., Ramírez-Portilla, C., Sánchez, J.A., Rodríguez, E., McFadden, C.S., 2018. Universal target-enrichment baits for anthozoan (Cnidaria) phylogenomics: New approaches to long-standing problems. Mol. Ecol. Resour. 18, 281–295. https://doi.org/10.1111/1755-0998.12736

41. Quek, R.Z.B., Jain, S.S., Neo, M.L., Rouse, G.W., Huang, D., 2020. Transcriptome-based target-enrichment baits for stony corals (Cnidaria: Anthozoa: Scleractinia). Mol. Ecol. Resour. 1755–0998.13150. https://doi.org/10.1111/1755-0998.13150

42. Rabosky, D.L., 2015. No substitute for real data: A cautionary note on the use of phylogenies from birth-death polytomy resolvers for downstream comparative analyses. Evolution (N. Y). 3207–3216. https://doi.org/10.1111/evo.12817

43. Rabosky, D.L., Chang, J., Title, P.O., Cowman, P.F., Sallan, L., Friedman, M., Kaschner, K., Garilao, C., Near, T.J., Coll, M., Alfaro, M.E., 2018. An inverse latitudinal gradient in speciation rate for marine fishes. Nature 559, 392–395. https://doi.org/10.1038/s41586-018-0273-1

44. Renema, W., Pandolfi, J.M., Kiessling, W., Bosellini, F.R., Klaus, J.S., Korpanty, C., Rosen, B.R., Santodomingo, N., Wallace, C.C., Webster, J.M., Johnson, K.G., 2016. Are coral reefs victims of their own past success ? 1–7.

45. Richards, Z.T., Berry, O., van Oppen, M.J.H., 2016. Cryptic genetic divergence within threatened species of Acropora coral from the Indian and Pacific Oceans. Conserv. Genet. 17, 577–591. https://doi.org/10.1007/s10592-015-0807-0

46. Richards, Z.T., Miller, D.J., Wallace, C.C., 2013. Molecular phylogenetics of geographically restricted Acropora species: Implications for threatened species conservation. Mol. Phylogenet. Evol. 69, 837–851. https://doi.org/10.1016/j.ympev.2013.06.020

47. Romano, S.L., Palumbi, S.R., 1996. Evolution of Scleractinian Corals Inferred from Molecular Systematics. Science (80-.). 271, 640–642. https://doi.org/10.1126/science.271.5249.640

48. Schmidt-Roach, S., Miller, K.J., Andreakis, N., 2013. Pocillopora aliciae: A new species of scleractinian coral (Scleractinia, Pocilloporidae) from subtropical Eastern Australia. Zootaxa. https://doi.org/10.11646/zootaxa.3626.4.11

49. Shearer, T.L., Van Oppen, M.J.H., Romano, S.L., Wörheide, G., 2002. Slow mitochondrial DNA sequence evolution in the Anthozoa (Cnidaria). Mol. Ecol. 11, 2475–2487. https://doi.org/10.1046/j.1365-294X.2002.01652.x

50. Sigovini, M., Keppel, E., Tagliapietra, D., 2016. Open Nomenclature in the biodiversity era. Methods Ecol. Evol. 7, 1217–1225. https://doi.org/10.1111/2041-210X.12594

51. Smith, B.T., Harvey, M.G., Faircloth, B.C., Glenn, T.C., Brumfield, R.T., 2014. Target capture and massively parallel sequencing of ultraconserved elements for comparative studies at shallow evolutionary time scales. Syst. Biol. 63, 83–95. https://doi.org/10.1093/sysbio/syt061

52. Stamatakis, A., 2014. RAxML version 8: A tool for phylogenetic analysis and post-analysis of large phylogenies. Bioinformatics 30, 1312–1313. https://doi.org/10.1093/bioinformatics/btu033

53. Terraneo, T.I., Arrigoni, R., Benzoni, F., Tietbohl, M.D., Berumen, M.L., 2017. Exploring the genetic diversity of shallow-water Agariciidae (Cnidaria: Anthozoa) from the Saudi Arabian Red Sea. Mar. Biodivers. 47, 1065–1078. https://doi.org/10.1007/s12526-017-0722-3

54. Thomas, G.H., Hartmann, K., Jetz, W., Joy, J.B., Mimoto, A., Mooers, A.O., 2013.x PASTIS: an R package to facilitate phylogenetic assembly with soft taxonomic inferences. Methods Ecol. Evol. 4, 1011–1017. https://doi.org/10.1111/2041-210X.12117

55. Thomson, S.A., Pyle, R.L., Ahyong, S.T., Alonso-Zarazaga, M., Ammirati, J., Araya, J.F., Ascher, J.S., Audisio, T.L., Azevedo-Santos, V.M., Bailly, N., Baker, W.J., Balke, M., Barclay, M.V.L., Barrett, R.L., Benine, R.C., Bickerstaff, J.R.M., Bouchard, P., Bour, R., Bourgoin, T., Boyko, C.B., Breure, A.S.H., Brothers, D.J., Byng, J.W., Campbell, D., Ceríaco, L.M.P., Cernák, I., Cerretti, P., Chang, C.-H., Cho, S., Copus, J.M., Costello, M.J., Cseh, A., Csuzdi, C., Culham, A., D’Elía, G., d’Udekem d’Acoz, C., Daneliya, M.E., Dekker, R., Dickinson, E.C., Dickinson, T.A., van Dijk, P.P., Dijkstra, K.-D.B., Dima, B., Dmitriev, D.A., Duistermaat, L., Dumbacher, J.P., Eiserhardt, W.L., Ekrem, T., Evenhuis, N.L., Faille, A., Fernández-Triana, J.L., Fiesler, E., Fishbein, M., Fordham, B.G., Freitas, A.V.L., Friol, N.R., Fritz, U., Frøslev, T., Funk, V.A., Gaimari, S.D., Garbino, G.S.T., Garraffoni, A.R.S., Geml, J., Gill, A.C., Gray, A., Grazziotin, F.G., Greenslade, P., Gutiérrez, E.E., Harvey, M.S., Hazevoet, C.J., He, K., He, X., Helfer, S., Helgen, K.M., van Heteren, A.H., Hita Garcia, F., Holstein, N., Horváth, M.K., Hovenkamp, P.H., Hwang, W.S., Hyvönen, J., Islam, M.B., Iverson, J.B., Ivie, M.A., Jaafar, Z., Jackson, M.D., Jayat, J.P., Johnson, N.F., Kaiser, H., Klitgård, B.B., Knapp, D.G., Kojima, J., Kõljalg, U., Kontschán, J., Krell, F.-T., Krisai-Greilhuber, I., Kullander, S., Latella, L., Lattke, J.E., Lencioni, V., Lewis, G.P., Lhano, M.G., Lujan, N.K., Luksenburg, J.A., Mariaux, J., Marinho-Filho, J., Marshall, C.J., Mate, J.F., McDonough, M.M., Michel, E., Miranda, V.F.O., Mitroiu, M.-D., Molinari, J., Monks, S., Moore, A.J., Moratelli, R., Murányi, D., Nakano, T., Nikolaeva, S., Noyes, J., Ohl, M., Oleas, N.H., Orrell, T., Páll-Gergely, B., Pape, T., Papp, V., Parenti, L.R., Patterson, D., Pavlinov, I.Y., Pine, R.H., Poczai, P., Prado, J., Prathapan, D., Rabeler, R.K., Randall, J.E., Rheindt, F.E., Rhodin, A.G.J., Rodríguez, S.M., Rogers, D.C., Roque, F. de O., Rowe, K.C., Ruedas, L.A., Salazar-Bravo, J., Salvador, R.B., Sangster, G., Sarmiento, C.E., Schigel, D.S., Schmidt, S., Schueler, F.W., Segers, H., Snow, N., Souza-Dias, P.G.B., Stals, R., Stenroos, S., Stone, R.D., Sturm, C.F., Štys, P., Teta, P., Thomas, D.C., Timm, R.M., Tindall, B.J., Todd, J.A., Triebel, D., Valdecasas, A.G., Vizzini, A., Vorontsova, M.S., de Vos, J.M., Wagner, P., Watling, L., Weakley, A., Welter-Schultes, F., Whitmore, D., Wilding, N., Will, K., Williams, J., Wilson, K., Winston, J.E., Wüster, W., Yanega, D., Yeates, D.K., Zaher, H., Zhang, G., Zhang, Z.-Q., Zhou, H.-Z., 2018. Taxonomy based on science is necessary for global conservation. PLOS Biol. 16, e2005075. https://doi.org/10.1371/journal.pbio.2005075

56. Tonini, J.F.R., Beard, K.H., Ferreira, R.B., Jetz, W., Pyron, R.A., 2016. Fully-sampled phylogenies of squamates reveal evolutionary patterns in threat status. Biol. Conserv. https://doi.org/10.1016/j.biocon.2016.03.039

57. Upham, N., Esselstyn, J., Jetz, W., 2019. Inferring the mammal tree: species-level sets of phylogenies for questions in ecology, evolution, and conservation. PLoS Biol. 1–44.

58. Van Oppen, M.J.H., Willis, B.L., Miller, D.J., 1999. Atypically low rate of cytochrome b evolution in the scleractinian coral genus Acropora. Proc. R. Soc. B Biol. Sci. 266, 179– 183. https://doi.org/10.1098/rspb.1999.0619

59. Veron, J.E.N., Wallace, C.C., 1984. Scleractinia of Eastern Australia, Part 5: Acroporidae. Aust Inst Mar Sci Monogr Ser 6, 485.

60. Wallace, C.C., 1999. Staghorn Corals of the World, Staghorn Corals of the World. CSIRO Publishing. https://doi.org/10.1071/9780643101388

61. Wallace, C.C., Done, B.J., Muir, P.R., 2012. Revision and catalogue of worldwide staghorn corals Acropora and Isopora (Scleractinia: Acroporidae) in the Museum of Tropical Queensland. Australia: Memoirs of the Queensland Museum.

62. Ying, H., Cooke, I., Sprungala, S., Wang, W., Hayward, D.C., Tang, Y., Huttley, G., Ball, E.E., Forêt, S., Miller, D.J., 2018. Comparative genomics reveals the distinct evolutionary trajectories of the robust and complex coral lineages. Genome Biol. 19, 175. https://doi.org/10.1186/s13059-018-1552-8

