## Supplementary material for "An enhanced target-enrichment bait set for Hexacorallia provides phylogenomic resolution of the staghorn corals (Acroporidae) and close relatives": Suppl Figs

### Supplemental Figures

**Figure S1.** RAxML cladograms for UCE only vs exon only 75% complete matrix alignments. Internal nodes were rotated using the *cophylo()* function in the R package phytools (Revell, 2012) to maximise comparison among topologies. Bootstrap values of 100% (red) and great than or equal to 80% (blue) are indicated at internal nodes.

**Figure S2.** RAxML cladograms for UCE only vs exon only 95% complete matrix alignments. Internal nodes were rotated using the *cophylo()* function in the R package phytools (Revell, 2012) to maximise comparison among topologies. Bootstrap values of 100% (red) and great than or equal to 80% (blue) are indicated at internal nodes.

**Figure S3.** RAxML cladograms for combined UCE/exon 75% vs combined UCE/exon 95% complete matrix alignments. Internal nodes were rotated using the *cophylo()* function in the R package phytools (Revell, 2012) to maximise comparison among topologies. Bootstrap values of 100% (red) and great than or equal to 80% (blue) are indicated at internal nodes.

**Figure S4.** Comparison of RAxML cladogram and ExaBayes cladogram produced from the combine UCE/exon 95% complete matrix alignment. Internal nodes were rotated using the *cophylo()* function in the R package phytools (Revell, 2012) to maximise comparison among topologies. RAxML Bootstrap values of 100% (red) and great than or equal to 80% (blue) are indicated at internal nodes on RAxML tree (right), while Bayesian posterior probability values of 1 (red) or greater than or equal to 0.7 (blue) are indicated on the ExaBayes tree (right).

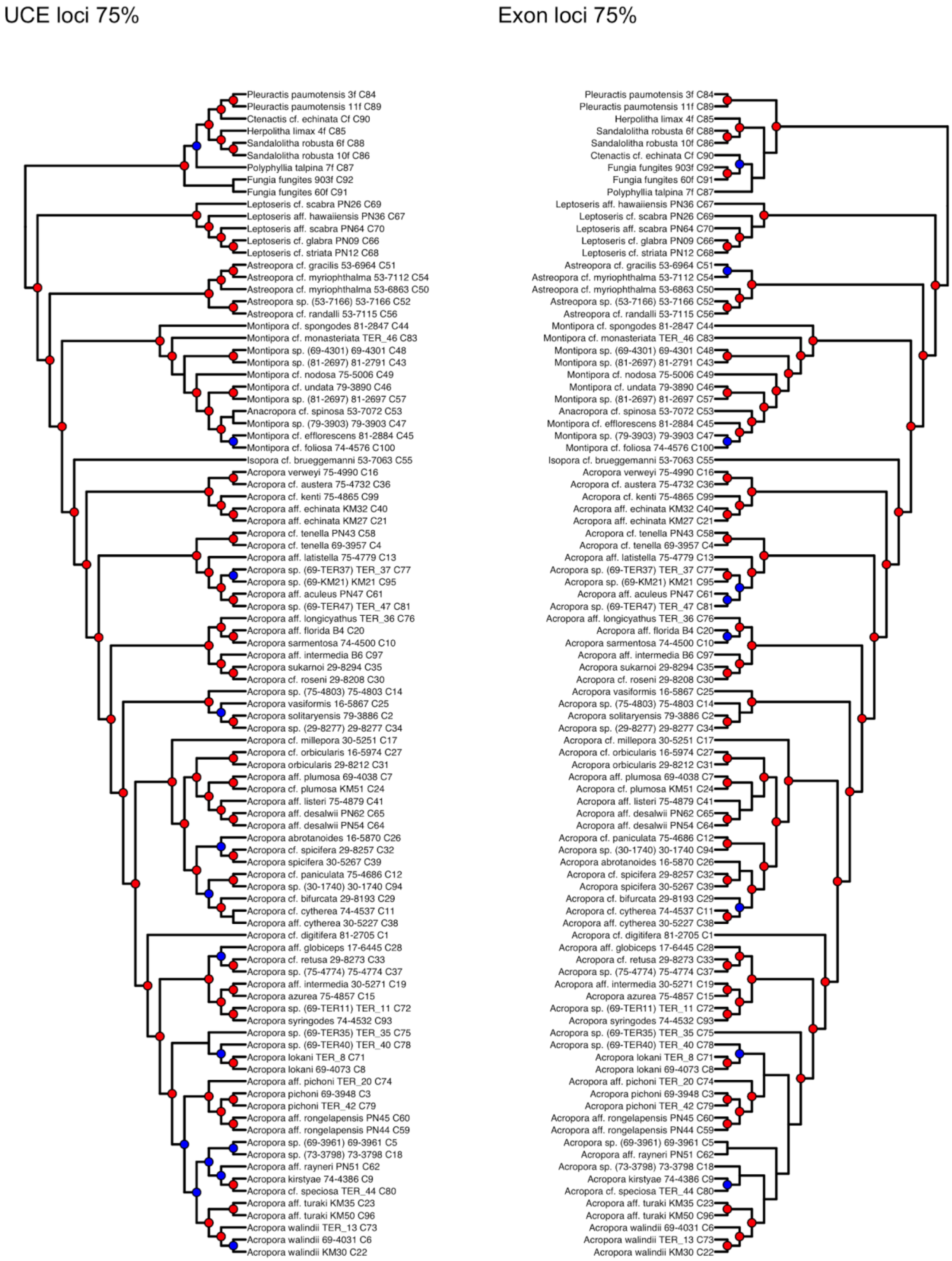


**Figure S1.** RAxML cladograms for UCE only vs exon only 75% complete matrix alignments.

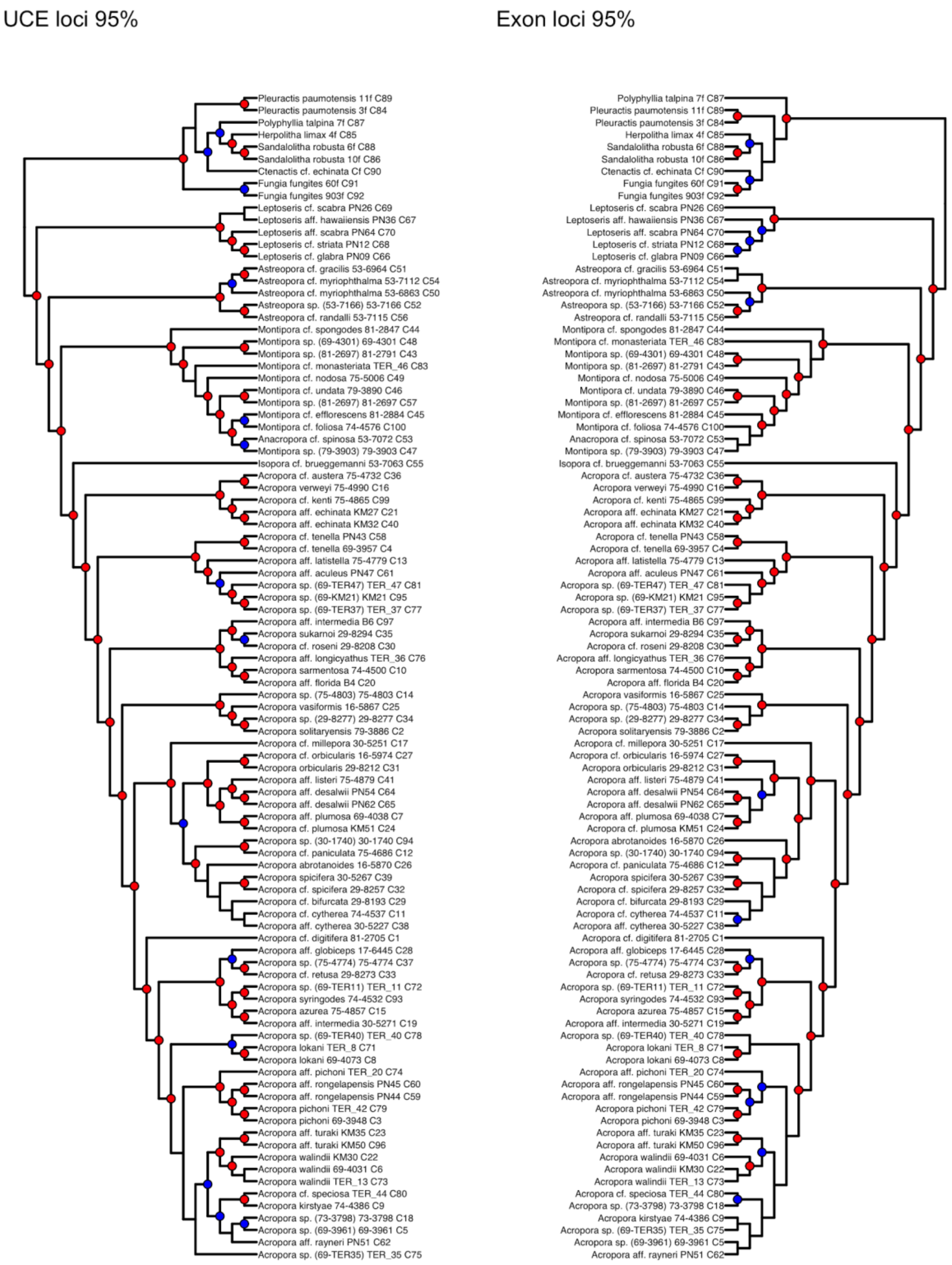


**Figure S2.** RAxML cladograms for UCE only vs exon only 95% complete matrix alignments.

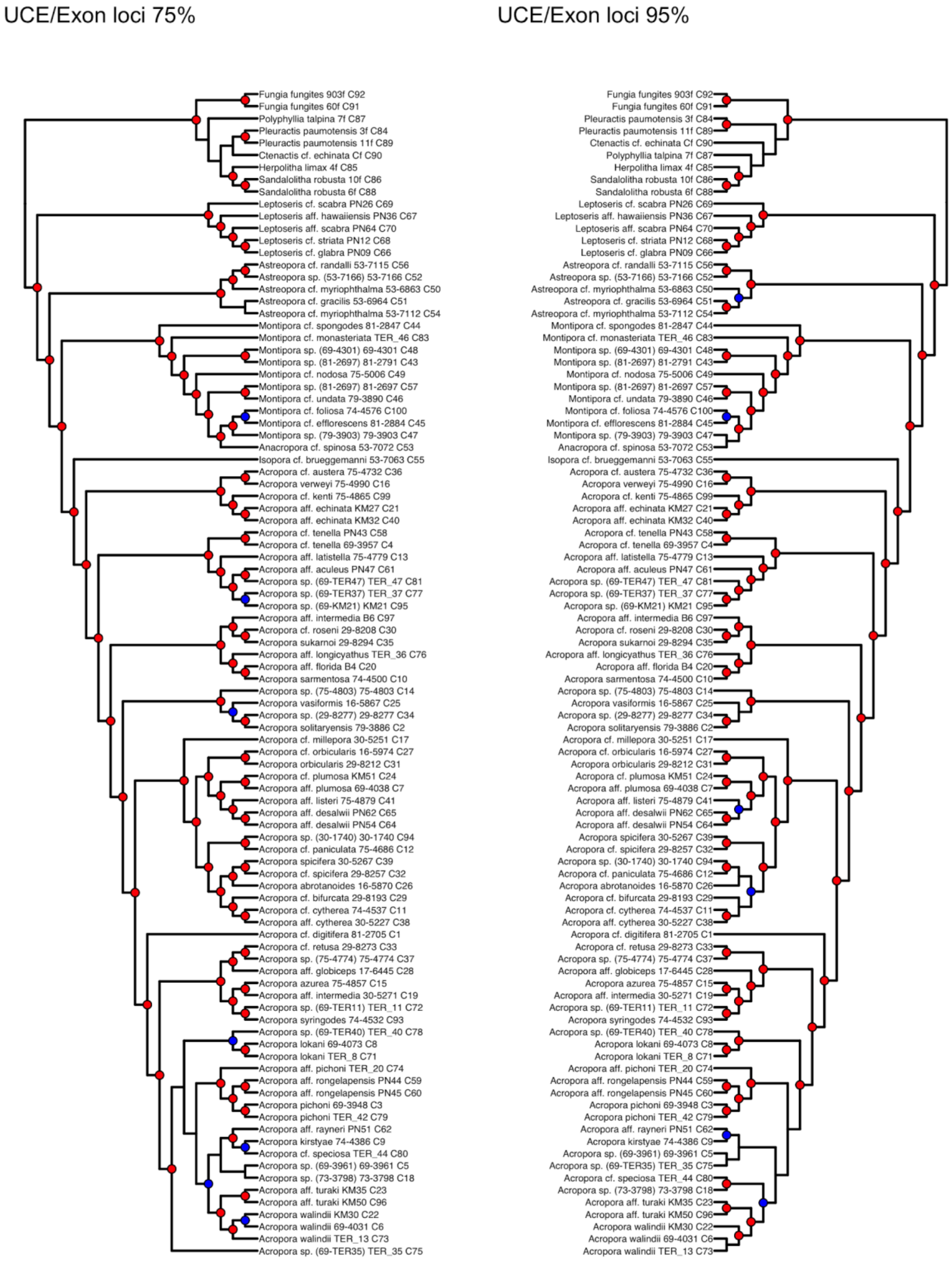


**Figure S3.** RAxML cladograms for combined UCE/exon 75% vs combined UCE/exon 95% complete matrix alignments.

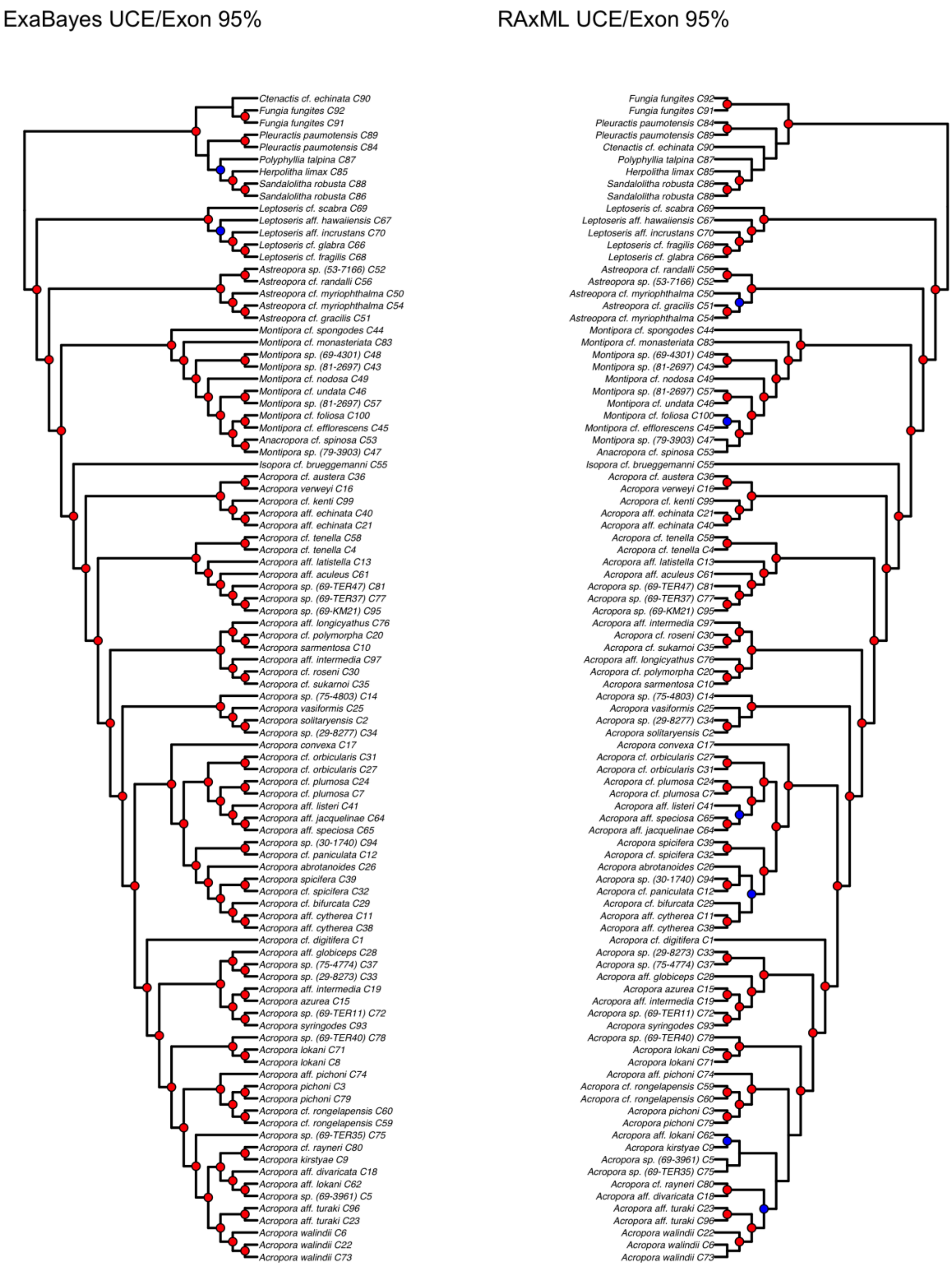


**Figure S4.** Comparison of RAxML cladogram and ExaBayes cladogram produced from the combine UCE/exon 95% complete matrix alignment.

### References

Revell, L. J. (2012). phytools: an R package for phylogenetic comparative biology (and other things). *Methods in Ecology and Evolution*, *3*(2), 217–223. doi:10.1111/j.2041-210X.2011.00169.x
